# Mosaic human cortical organoids model mTOR-related focal cortical dysplasia through DEPDC5 loss-of-function

**DOI:** 10.1101/2025.06.25.661323

**Authors:** Marina Maletic, Sara Bizzotto, Théo Ribierre, Kenza Guerdoud, Corentin Raoux, Marion Doladilhe, Carine Dalle, Fabienne Picard, Stéphanie Baulac

**Affiliations:** Sorbonne Université, Paris Brain Institute (ICM), Inserm, CNRS, AP-HP, Pitié-Salpêtrière Hospital, Paris, France; Université Paris Cité, Imagine Institute, Paris, France; Department of Basic Neurosciences, University of Geneva, Geneva, Switzerland; NeuroNA Human Cellular Neuroscience Platform, Fondation Campus Biotech Geneva, Geneva, Switzerland; Department of Clinical Neurosciences, University Hospitals and Medical School of Geneva, Geneva, Switzerland

## Abstract

Focal cortical dysplasia type II (FCDII), a leading cause of pediatric drug-resistant focal epilepsy, results from brain somatic variants in genes of the mTOR pathway, including germline and somatic second-hit loss-of-function variants in the mTOR repressor *DEPDC5*. Here, we investigated the effects of mosaic *DEPDC5* two-hit variants on cortical development and neuronal activity using patient-derived human cortical organoids (hCOs). Mosaic hCOs displayed increased mTOR activity and altered neural rosette densities, which were both rescued by treatment with the mTOR inhibitor rapamycin. In addition, mosaic hCOs presented dysmorphic-like neurons and increased neuronal excitability, recapitulating FCDII pathology. Longitudinal single-cell transcriptomics at three developmental stages revealed altered neuronal differentiation, dysregulated expression of genes associated with the Notch and Wnt pathways in neural progenitors, and of synaptic- and epilepsy-associated genes in excitatory neurons. We further identified cell-autonomous alterations in metabolism and translation in mosaic two-hit hCOs. This study provides novel insights into the consequences of mosaic biallelic *DEPDC5* deficiency on corticogenesis in the context of FCDII, highlighting both autonomous and non-cell autonomous effects.

## Introduction

Focal Cortical Dysplasia Type II (FCDII) is a neurodevelopmental disorder, part of the spectrum of cortical malformations. FCDII-associated epilepsy is typically resistant to antiseizure medications, necessitating neurosurgical resection of the epileptogenic zone for seizure control, and is the most prevalent cortical malformation in pediatric epilepsy surgeries^1^. FCDII can vary in extent, ranging from small, distinct cortical areas to an entire hemisphere, as seen in hemimegalencephaly (HME). The neuropathological hallmarks of FCDII and HME include cortical dyslamination and the presence of dysmorphic neurons (FCDIIa type), and in some cases, balloon cells (FCDIIb type)^2^. Dysmorphic neurons, characterized by an enlarged soma and an accumulation of neurofilaments, are thought to play a critical role in contributing to epileptic discharge generation, due to their hyperexcitability and presence within the epileptogenic focus^3^.

Recent studies have revealed the contribution of somatic mosaicism to various neurodevelopmental conditions due to the acquisition of postzygotic (i.e., somatic variants) during brain development^4,5^. Notably, pathogenic somatic variants in genes of the mTOR (mechanistic target of rapamycin) pathway have been increasingly recognized as a major cause of FCDII^6^. In FCDII, somatic variants are thought to arise during corticogenesis in dorsal pallium progenitors lining the ventricular zone^7^, but recent single-cell genomic and transcriptomic analyses reveal that mutations are present across multiple brain cell types, suggesting they may originate at earlier developmental stages^8^. The level of mosaicism (percentage of cells in the tissue that carry the variant), reflected by the variant allele frequency (VAF), can vary widely, being as low as ∼1% in FCDII and up to 30% in HME^9^.

Deep sequencing of surgical FCDII brain tissue has revealed the contribution of somatic gain-of-function (GoF) heterozygous variants in genes coding for PI3K-mTOR pathway activators such as *PIK3CA*, *AKT3* and *MTOR*. Furthermore, two-hit loss-of-function (LoF) variants – germline and somatic – have been found in genes coding for repressors of the PI3K-mTOR pathway, such as *DEPDC5* or *TSC1/TSC2*^10–14^. All these variants cause hyperactivation of the mTOR pathway, which in turn may dysregulate cell growth, differentiation, metabolism, and function.

DEPDC5 is a key component of the GATOR1 (Gap Activity TOward Rags 1) complex that represses mTORC1 signaling in response to amino acids (leucine and arginine specifically)^15^. *DEPDC5* mutations are major contributors to both lesional and non-lesional focal epilepsies^16–18^. A recent study identified *DEPDC5* as the most significant gene in non-acquired focal epilepsy among 9,219 cases^19^. Moreover, *DEPDC5* variants account for 13% of FCDII or HME cases^20^. Notably, *DEPDC5* variants are predominantly associated with FCDIIa, rather than FCDIIb, and do not generate balloon cells^21^. Several studies support Knudson’s two-hit mechanism in FCDII pathogenesis to explain the focal and mosaic nature of FCDII lesions, where a somatic second-hit occurs in addition to the germline *DEPDC5* variant in brain tissue^11,12,22–25^. We and others have shown that these somatic variants are enriched in dysmorphic neurons^11,25^. However, key questions remain regarding the relative contributions of heterozygous versus biallelic inactivation to epileptogenesis and FCDII pathogenesis.

In this study, we examined how mosaic *DEPDC5* biallelic inactivation affects cortical development and how it contributes to epileptogenesis and FCDII insurgence by generating patient-derived heterozygous (Het) and mosaic two-hit (Mos) human cortical organoids (hCOs). For this purpose, we employed longitudinal single-cell RNA-sequencing (scRNA-seq) across three developmental stages, as well as electrophysiological recording. Our findings reveal that Het and Mos hCOs display variable neurodevelopmental, transcriptomic, and functional alterations. Notably, however, only Mos hCOs recapitulated key FCDII characteristics, including cytomegalic mTOR-hyperactive neurons and neuronal network hyperactivity. Furthermore, our data highlight both cell-autonomous and non-cell-autonomous effects of *DEPDC5* loss during cortical development.

## Materials and Methods

### Generation and Characterization of hiPSC lines

Patient-derived hiPSCs were generated from peripheral blood mononuclear cells isolated from two male subjects^41^ using viral-free episomal reprogramming (Phenocell). Lines fulfilled standard reprogramming criteria, including serology, mycoplasma testing, embryoid body formation, pluripotency marker expression, and genomic stability. The study received ethical approval (Inserm N° C1456 GENEPI) with informed consent.

### CRISPR-Cas9 Gene Editing

A DEPDC5-/- line was generated by introducing p.L1420Ffs*154 mutation into the second allele of patient DEPDC5+/- hiPSCs using CRISPR-Cas9-mediated homologous repair. A CAG-EGFP-pA construct was inserted at the h*TIGRE* locus by CRISPR-Cas9. CRISPR reagents were delivered via electroporation (Neon system, Invitrogen). Two isogenic DEPDC5-/- clones were validated based on EGFP expression and normal karyotype (**Figure S1A and S1B**). Both isogenic *DEPDC5*^-/-^1 and *DEPDC5*^-/-^2 clones were used interchangeably in the study and are referred to as *DEPDC5*^-/-^. The isogenic control line was generated by targeting the mutated allele with the corrected sequence. All hiPSC lines will be registered in the hPSCreg database (https://hpscreg.eu/).

### hiPSC Culture

hiPSC lines were maintained in mTESR1 medium on laminin-521 or Vitronectin XF-coated dishes at 37°C, 5% CO2, passaged at 80% confluency using Accutase or ReLeSR (Stem Cell Technologies), and tested monthly for mycoplasma.

### Human Cortical Organoid Generation

hiPSCs were differentiated into human cortical organoids (hCOs) following a directed protocol towards the dorsal cortical fate^26^, with adaptations: DMSO omitted at day -2, mTeSR1 medium used instead of Essential 8, and NeuroCult1 supplement (vitamin A-free) instead of B27 (**Figure S1D**). To initiate differentiation, hiPSCs were dissociated into single-cell suspensions using Accutase. 4 × 10^6 cells/line were seeded in 24-well AggreWell 800 plates in mTeSR1 medium supplemented with 10 µM Rock inhibitor at 5% CO_2_ and 37°C for 24 hours to form embryoid bodies. For mosaic organoids, EGFP+ DEPDC5-/- and EGFP- DEPDC5+/- cells were combined at 50% (or 15% ratios. Embryoid bodies were transferred to TeSR-E6 medium containing Dorsomorphin (2.5 µM, SB-431542(10 µM) and XAV-939 (2.5 µM). Neural induction used Neurobasal-A (Life Technologies) supplemented with FGF2/EGF (20 ng/ml, days 6-24), then BDNF/NT-3 (20 ng/ml, days 25-42). From day 43, hCOs were maintained in Neurobasal-A with penicillin-streptomycin (100 U/ml). Medium was changed daily until day 14, then every other day, with orbital shaking from day 30.

For amino acid deprivation, 6-month hCOs were incubated for 4 hours in leucine/arginine-deficient SILAC Neurobasal Medium (Athena Enzyme Solutions) before immediate harvest.

For mTOR inhibition, hCOs received 20nM rapamycin or vehicle (ethanol) in Neurobasal medium from day 25 until analysis, as previously described^35^.

### Western blotting

Samples were lysed with protease and phosphatase inhibitors, centrifuged at 14,000 rpm, and supernatants were stored at -80°C. Protein concentrations were determined by the BCA protein assay. Equal protein amounts were separated onto 4-12% NuPAGE 10-well gels, blotted, and membranes were blocked in PBS with 0.1% Tween-20 and 3% BSA, before overnight primary antibody incubation at 4°C, followed by HRP-coupled secondary antibody. Protein bands were visualized with ECL peroxidase substrates, quantified with FIJI software and normalized to β-actin. Five hCOs were pooled per genotype and per experiment, with five technical replicates.

### Immunostaining

hCOs were fixed in 4% PFA, cryopreserved in 30% sucrose, embedded in O.C.T., and frozen in dry ice. 20 µm sections were blocked (PBS, 0.4% Triton X-100, 10% goat serum), incubated with primary antibodies overnight, then with secondary antibodies and DAPI for 2 hours. Slides were mounted with Fluoromount-G and imaged on Zeiss Apotome or Nikon confocal microscopes. Each experiment was repeated in at least three independent organoid sections.

For 3D immunostaining, whole fixed hCOs underwent serial detergent permeabilization steps, blocking, and prolonged antibody incubations with continuous shaking. hCOs were clarified with RapiClear before confocal imaging. SATB2 protein expression was quantified using ImageJ’s analyze particles function to measure SATB2+ particle area relative to total slice area.

### Antibodies

Primary antibodies used included: β-Actin (Cell Signaling Technology, Cat# 4970; Sigma-Aldrich, Cat# A2066), DEPDC5 (Abcam, Cat# ab213181, ab185565), DLK1 (Abcam, Cat# ab21682), BLBP (Abcam, Cat# ab32423), GFP (Aves Labs, Cat# GFP-1020; Invitrogen, Cat# A-6455), NeuN (Millipore, Cat# MAB377, ABN78), phospho-S6 ribosomal protein (Ser240/244) (Cell Signaling Technology, Cat# 5364), SATB2 (Abcam, Cat# ab51502), neurofilament SMI311 (BioLegend, Cat# BLE837801), SOX2 (Thermo Fisher Scientific, Cat# 14-9811-82; Millipore, Cat# AB5603), and ZO-1 (Thermo Fisher Scientific, Cat# 33-9100; Cell Signaling Technology, Cat# 13663S). Secondary antibody: anti-rabbit IgG HRP-linked (Cell Signaling Technology, Cat# 7074).

### Multi-Electrode Array (MEA) recordings

Electrophysiological measurements were conducted using a 64-channel MEA system (MED64, Alpha MED Scientific), with poly-L-ornithine/laminin-521-coated probes (in neurobasal medium at 37°C, 48 hours post-medium change. Extracellular field potentials were filtered (100 Hz-10 kHz band-pass filter) and sampled at 20 kHz. Spike detection was performed using MOBIUS software with a detection threshold set at 6x the standard deviation of noise. Data analysis was performed using a custom R script available at https://gitlab.com/icm-institute/dac/biostats/MEASpikeR. Electrodes with ≥5 spikes/minute were considered active. Firing rate was calculated as the total spike number per electrode over 3 minutes.

### Single-cell RNA sequencing (scRNA-seq)

<u>Sample preparation.</u> hCOs were prepared for scRNA-seq following a previously reported protocol^42^. Five hCOs were harvested per library with the following batches per condition: CT (1 month N=1, 3 months N=1, 6 months N=2); Het (1 month N=2, 3 months N=1, 6 months N=1); Mos (1 month N=2, 3 months N=1, 6 months N=2). Dissociated cells were resuspended in ice-cold solution (0.1% BSA and 25 mM HEPES in DPBS), incubated with eBioscience Fixable Viability Dye eFluor 660 65-0864 (1:1000) for 30 minutes, filtered (50 µm) and sorted (BioRad S3e). Viability gating used eFluor660 dye, EGFP gating used the intrinsic EGFP signal. Viable cells were collected in BSA-coated tubes, counted and checked for viability (≥80% required). Processing used 10X Genomics Chromium Next GEM Single Cell 3C Kit v3.1. Approximately 20,000 cells were loaded per library to retrieve ∼10,000 cells, with the same-day GEM generation, barcoding, and cDNA amplification. Libraries were sequenced on Illumina NovaSeq6000 targeting ∼50,000 read-pairs per cell.

### scRNA-seq data processing

Base call files (BCLs) from Illumina NGS raw data were demultiplexed into FASTQ files using 10X Genomics cellranger mkfastq. Alignment to hg38 customized with the *EGFP* gene sequence, and barcode and unique molecular identifier (UMI) counting for generation of feature/barcode count matrices were done with cellranger count. Filtered count matrices were used using Seurat (version 4.1.0)^43^. Further filters were applied to keep: 1) genes expressed in ≥5 cells; 2) cells expressing ≥ 200 genes; 3) cells with < 8% mitochondrial RNA; 4) cells with <70% ribosomal RNA; 5) cells with 500-20,000 UMIs. Additional filters included the removal of all cells found before the local minimum based on the function *(f): nFeature -> density(nFeature)*, with *nFeature* representing the number of features arranged in ascending order. DoubletFinder^44^ removed doublets (<10% per sample). Since we did not observe significant batch effects on our data, integration was unnecessary; libraries were normalized using *NormalizeData* with the *LogNormalize* method, and by regressing out mitochondrial percentage and UMI count. Variable features were identified using *FindVariableFeatures* function. Data were further corrected to reduce the effect of cell cycle. Data scaling was done with *ScaleData* and dimensional reduction used *RunPCA* and *RunUMAP*.

### Cell type annotation

Clusters were annotated in CT hCO data using canonical cortical cell markers^27,45^ (Figure S2A). To verify cluster annotation, label transfer using a reference atlas was performed using SingleR^46^. For label transfer, we used temporally matched reference data: 23 days-1.5 months reference for 1-month hCOs, 2-4 months reference for 3-month hCOs, and 4-6 months reference for 6-month hCOs. Het and Mos hCO clusters were subsequently annotated by label transfer using Seurat *FindTransferAnchors* followed by *TransferData* using CT hCOs data as reference (Figure S2B), and further refining the annotation based on canonical cell type marker expression.

### RNA velocity and trajectory analysis

For RNA velocity analyses, each genotype was analyzed separately (pooled timepoints). RNA velocities, latent time, and driver genes were calculated using scVelo’s dynamic model^47^.

### Differential gene expression analysis

Differential gene expression used Seurat *FindMarkers* function with Wilcoxon Rank Sum test (minimum 1% cell percentage, logfc threshold at 0.1). Only genes with adjusted <0.05 (Bonferroni) were retained.

### Gene ontology analyses

For gene ontology (GO) biological process enrichment analysis, we used ClusterProfiler’s enrichGO function (v.4.2.2) ^48^, with the following parameters: 1) *p*-value < 0.01; 2) *q*-value < 0.1; 3) gene size at 3-1000; 4) adjusted *p*-values calculated with the Benjamini & Hochberg correction. We set as background (universe) all genes expressed across hCOs genotypes and timepoints. We used synaptic gene ontology (SynGO) with the DEG lists obtained for each comparison as input. We focused the enrichment analysis on biological processes.

### Statistical analyses

Statistical analyses of imaging, Western Blot and MEA data were performed using the GraphPad Prism (v. 9.4.1). Values are presented as mean ± standard deviation, with specific tests and sample sizes detailed in figure legends. The selection of hCOs for amino acid deprivation and rapamycin treatment was randomized. All samples were included in the analysis to ensure comprehensive and unbiased results. Significance thresholds: *, *p* < 0.05; **, *p* < 0.01; ***, *p* < 0.001; ****, *p* < 0.0001.

### Data and code availability

Single-cell RNA-sequencing data will be available at the European Genome-phenome Archive (EGA). No original computational tools were generated for this study.

## Results

### *DEPDC5* mosaic hCOs display mTOR-hyperactive dysmorphic-like neurons

To investigate how *DEPDC5* mutation alters early cortical development and model FCDII pathogenesis, we derived human induced pluripotent stem cells (hiPSCs) from two brothers: a patient with focal epilepsy and FCDII carrying a heterozygous loss-of-function *DEPDC5* variant (p.L1420Ffs*154), and his unaffected, non-carrier sibling who serve as an age- and sex-matched control sharing 50% of the genetic background (**Figure 1A**). Using CRISPR-Cas9 editing, we then engineered an isogenic EGFP-tagged *DEPDC5* two-hit (homozygous) mutant line by introducing the same variant into the wildtype allele of the patient’s heterozygous hiPSCs (**Figure S1A and B** and **Materials and methods**). An isogenic control hiPSC line was also generated by CRISPR-Cas9 (*DEPDC5*^+/+^) but was withdrawn from most experiments since its quality was questionable as we repeatedly observed a tendency to spontaneously differentiate. Therefore, we used three distinct hiPSC lines in this study: 1) a *DEPDC5*^+/-^ (EGFP-) line from the patient, 2) a control *DEPDC5*^+/+^ line from the asymptomatic brother, and 3) an isogenic *DEPDC5*^-/-^ (EGFP+) line. To validate the efficiency of gene-editing, we quantified the levels of DEPDC5 and EGFP proteins by Western blotting. We confirmed the absence of the DEPDC5 protein in the *DEPDC5*^-/-^ line, and half the amount of DEPDC5 in *DEPDC5*^+/-^ compared to *DEPDC5*^+/+^ (**Figure S1C**).

**Figure 1.**
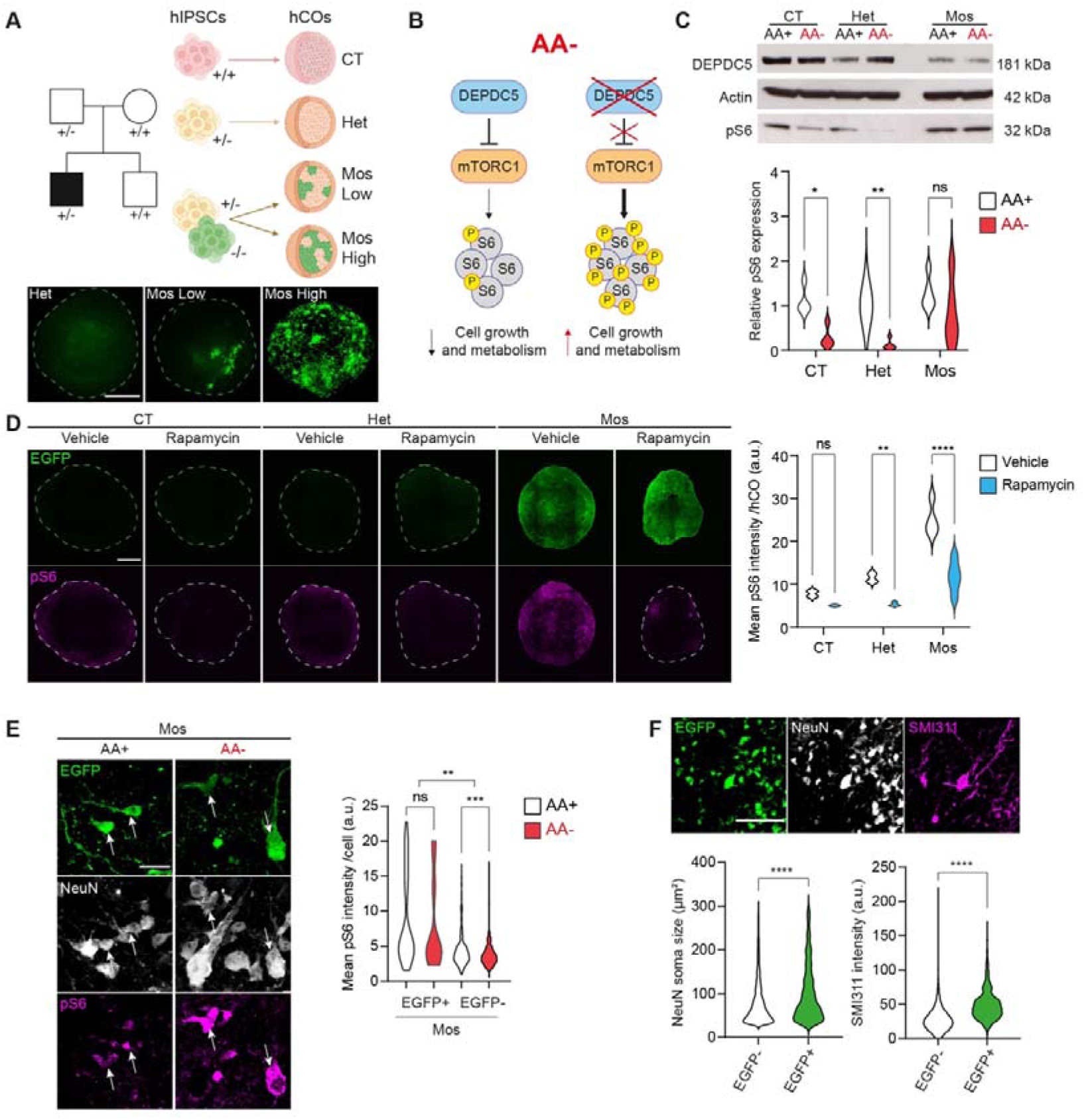
*DEPDC5* mosaic hCOs display dysmorphic-like neurons. (**A**) Left: Family pedigree showing inherited *DEPDC5* loss-of-function germline variant. HiPSCs were derived from the proband and unaffected sibling. Below: Endogenous EGFP signal in clarified 1-month Mos hCOs. Scale bar, 300µm. (**B**) Schematic of mTOR pathway activation (measured by S6 phosphorylation) under amino acid deprivation (AA-) with and without functional DEPDC5. (**C**) Western blot showing DEPDC5, pS6 and actin protein levels in 6-month hCOs under basal (AA+) and deprived (AA-) conditions. Quantification normalized to actin (n = 5 batches, 5 hCOs/genotype); Two-way ANOVA with Šidák’s correction. (**D**) pS6 immunofluorescence quantification in 3-month hCOs treated with vehicle or rapamycin (n = 4-5 hCOs/condition, 1 section/hCO). Scale bar, 1 mm. Two-way ANOVA with Šidák’s correction. (**E**) pS6 levels in EGFP- and EGFP+ NeuN+ neurons in 6-month Mos Low hCOs under AA+ and AA-conditions. Arrows: EGFP+/pS6+ neurons. Scale bar, 20 µm. n = 8 hCOs/condition from 2 batches; cells analyzed: EGFP+/AA+ = 33, EGFP+/AA- = 17, EGFP-/AA+ = 573, EGFP-/AA- = 535. Kruskal-Wallis test with Dunn’s correction. (**F**) Quantification of soma size and SMI311 intensity (normalized to soma area) in EGFP+ versus EGFP-NeuN+ neurons in 6-month Mos Low hCOs. Scale bar, 100 µm. n = 4 hCOs; cells analyzed: EGFP+ = 1,437, EGFP- = 6,045. Mann-Whitney test. *, *p* < 0.05; **, *p* < 0.01; ***, *p* < 0.001; ****, *p* < 0.0001.

The hiPSC lines were differentiated into dorsal cortical organoids following an established protocol^26^. We generated three types of human cortical organoids (hCOs): control (CT, *DEPDC5*^+/+^ cells), heterozygous (Het, *DEPDC5*^+/-^ EGFP-cells), and mosaic (Mos), the latter containing both *DEPDC5*^+/-^ EGFP- and *DEPDC5*^-/-^ EGFP+ cells, with either low (Mos Low, 15-25% EGFP+ cells) or high (Mos High, ∼50% EGFP+ cells) levels of mosaicism (**Figure 1A, Figure S1D** and **Materials and methods**. For simplicity, these three conditions (CT, Het, and Mos) are referred to as ‘genotypes’ throughout, although Mos hCOs contain cells of two distinct genotypes. Mos High hCOs were used for most experiments in the study, unless stated otherwise. Clarification and imaging of 1-month Mos hCOs, as well as flow cytometry analysis, confirmed the presence of EGFP+ cells in the expected proportions, with no evidence of positive or negative selection for the EGFP+ population (**Figure 1A, Figure S1E**). By day 14, only Mos hCOs were slightly larger than CT, while Iso and Het remained similar (**Figure S1F**). However, the isogenic control hCOs degraded rapidly, with their numbers decreasing significantly. This, along with the isogenic control hiPSC line’s tendency to differentiate spontaneously, indicated poor quality of the cell line, leading to its exclusion from downstream experiments. All other hCO genotypes (CT, Het and Mos) maintained structural integrity and characteristic neural rosette formation throughout 6 months of differentiation, indicating that *DEPDC5* deficiency does not significantly impair organoid development.

DEPDC5, within the GATOR1 complex, is a key repressor of mTOR activity in response to amino acid availability, especially leucine and arginine. In the absence of DEPDC5, mTOR signaling is predicted to remain constitutively active during amino acid deprivation (**Figure 1B**). To assess mTORC1 activity in mature 6-month hCOs, we stimulated the GATOR1 branch of the pathway by using a modified neurobasal medium deprived of leucine and arginine. Western blot analysis showed reduced phosphorylated S6 (pS6), an mTORC1 activity readout, in both CT and Het hCOs under amino acid deprivation (AA-). In contrast, Mos hCOs maintained pS6 levels comparable to baseline (AA+), indicating that GATOR1 failed to inhibit mTORC1 activity (**Figure 1C**). Phospho-S6 immunofluorescence analysis confirmed constitutive mTOR activation in Mos and, to a lesser extent, Het hCOs (**Figure S1G**). Administration of the mTORC1 inhibitor rapamycin significantly reduced pS6 levels in both Het and Mos hCOs, confirming that the constant pS6 levels resulted from impaired GATOR1-mediated mTORC1 inhibition (**Figure 1D**).

To determine the cell-autonomous effects of DEPDC5 loss, we analyzed pS6 immunofluorescence in NeuN-positive neurons from Low Mos hCOs, where sparse EGFP labeling enables clear distinction between DEPDC5^-/-^ (EGFP+) and DEPDC5^+/-^ (EGFP-) cells. EGFP+ neurons exhibited significantly higher basal pS6 levels compared to neighboring EGFP-neurons within the same Mos hCO and maintained these levels during amino acid deprivation, indicating constitutive mTORC1 activation. In contrast, EGFP-neurons showed decreased pS6 levels under amino acid-deprived conditions (**Figure 1E**).

We searched for the presence of FCDII hallmark dysmorphic neurons by performing co-immunostaining of EGFP, NeuN and SMI311, a canonical neurofilament marker of dysmorphic neurons, in mature 6-month hCOs. EGFP+ (DEPDC5^-/-^) neurons in Mos hCOs exhibited enlarged soma size compared to EGFP- (DEPDC5^+/-^) neurons (26% increase; mean EGFP- = 78.59 μm^2^ ± 51.67, mean EGFP+ = 99.24 μm^2^ ± 64.47) and SMI311 accumulation (**Figure 1F**). Notably, no balloon-like cells were detected by vimentin staining, consistent with DEPDC5-related FCDII pathology, which typically presents with dysmorphic neurons but lacks balloon cells^9,21^.

These findings indicate that biallelic *DEPDC5* inactivation in hCOs leads to constitutive mTOR pathway activation, and is necessary to reproduce dysmorphic-like neurons, the main FCDII hallmark.

### *DEPDC5* loss-of-function leads to premature generation of upper-layer neurons

To investigate how DEPDC5 deficiency affects cortical development and cell fate acquisition, we performed single-cell RNA sequencing (scRNA-seq) of CT, Het and Mos hCOs at three key developmental stages: 1) progenitor expansion and initial neuronal production (1 month); 2) early-to-mid corticogenesis, when diverse cortical cell types start emerging (3 months); and 3) later corticogenesis (6 months). Following quality filtering (see **Materials and methods**), we analyzed a total of 73,685 cells for all three stages (28,637 CT; 14,972 Het; 30,076 Mos). Cell clusters were first annotated in CT hCOs at each timepoint based on the expression of canonical brain cell type marker genes (**Figure S2A**). To verify the annotation, we additionally performed label transfer from an organoid reference atlas of similar timepoints^27^ using SingleR. Subsequently, we annotated Het and Mos hCOs by label transfer, using CT as reference, and further verified and refined cluster annotations based on marker gene expression (**Figure S2B** and **Materials and methods**). In CT hCOs, we identified the expected cell types at each timepoint compared to similar studies^26–28^. At 1-month, CT hCOs were made primarily of progenitor cells, including FOXG1-negative progenitors and FOXG1-positive apical radial glia (aRG) expressing *HES1*. Furthermore, at 1 month, we detected FOXG1-negative neurons expressing *STMN2*, *GRIA2* and *SLC17A6*. In smaller proportions, we observed unspecified FOXG1-positive neurons (expressing *STMN2* and *GRIA2)*, preplate cells (expressing *EMX2*, *FOXP2* and *LHX9)*, and Cajal-Retzius-like cells (expressing *LHX5* and *RELN)*. At 3 months, we observed the appearance of *EOMES*-expressing intermediate progenitors (IP) and a proportion of lower-layer excitatory neurons (LL.Exc.N) expressing *NEUROD6*, *FEZF2* and *PDE1A*. By 6 months, in addition to aRG and IP, we detected outer RG (oRG) expressing *TNC*, *HOPX*, *FAM107A* and *MOXD1*. Concurrently, upper-layer excitatory neurons (UL.Exc.N) emerged, expressing *BHLHE22*, *PLXNA4*, *CUX2* and *SATB2*. At this time point, we also detected interneuron progenitors (IN.progenitors) expressing *BIRC5*, *GAD1/2*, *DLX1/2/5* and *SP8*, and inhibitory neurons (Inh.N) expressing *GAD1/2*, *DLX1/2/5*, *SP8, SCGN, PROX1, CALB2 and SLC32A1* (**Figure 2A, Figure S2A**).

**Figure 2.**
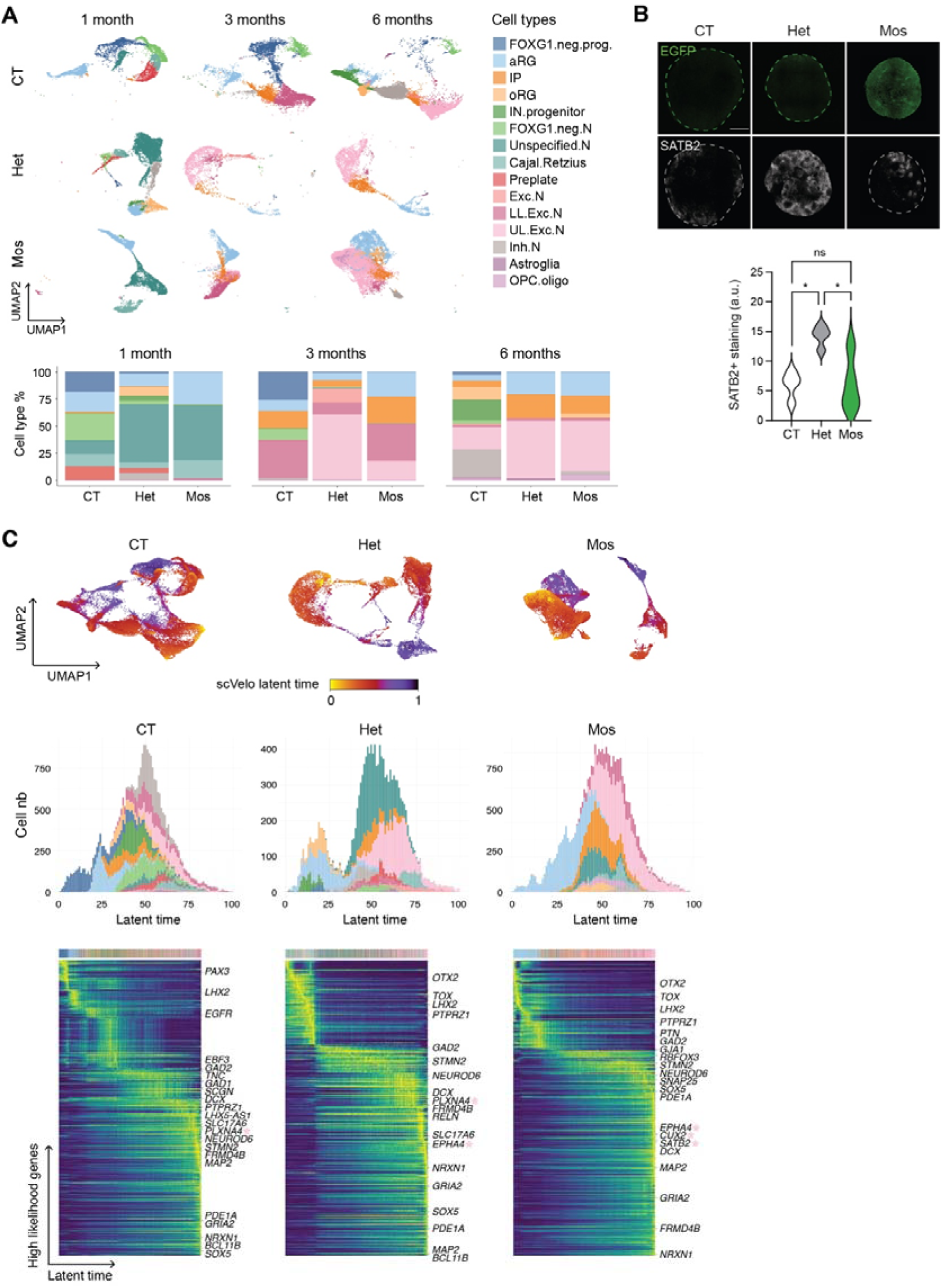
DEPDC5 deficiency alters differentiation trajectories. (**A**) Single-cell transcriptomic analysis of cortical organoids. Top: UMAP visualization of cell clusters from CT, Het, and Mos hCOs at 1, 3, and 6 months. Bottom: Quantification of cell type proportions across developmental timepoints. (**B**) SATB2 and EGFP immunofluorescence in 3-month whole hCOs. SATB2+ particle density quantified relative to section area (μm2). Scale bar: 500 µm. n = 4 hCOs/genotype (one batch), one section/hCO, one-way ANOVA with Tukey’s correction. (**C**) RNA velocity analysis across development. Top: UMAP visualization color-coded by scVelo latent time. Middle: Cell type distribution along latent time for each genotype. Bottom: Heatmaps showing expression dynamics of top 300 likelihood-ranked genes resolved along latent time. Colored stars indicate UL.Exc.N markers; other cell type marker genes are also annotated.

Het and Mos hCOs exhibited distinct cell type compositions compared to CT hCOs. This was observed as early as 1 month and persisted across all developmental stages. At 1 month, all clusters in Het and Mos hCOs expressed FOXG1 confirming telencephalic fate commitment, with Het already displaying a cluster expressing oRG markers. By 3 months, Mos and particularly Het hCOs showed high proportions of UL.Exc.N (18% and 60%, respectively), which were almost absent (<1%) in CT hCOs (**Figure 2A**). This was confirmed by SATB2 immunostaining, a marker of upper-layer neurons (**Figure 2B**). At 6 months, Het and Mos hCOs still exhibited higher proportions of UL.Exc.N compared to CT and a near absence of IN.progenitors and Inh.N (**Figure 2A**). To better explore cell type differentiation in the three genotypes, we performed RNA velocity analysis using scVelo (**Materials and methods**), which supported the genotype differences in cell fate acquisition. This analysis revealed distinct differentiation trajectories along latent time, reflected by genotype-specific transcriptional cascades defined by the expression dynamics of putative driver genes, including some neuronal subtype marker genes. Driver gene expression dynamics showed a more abrupt and possibly precocious transition from progenitors to more differentiated cell types in Het and Mos compared to CT, possibly underlying the premature production of UL.Exc.Ns. Coherent with this, Het and Mos hCOs displayed more UL.Exc.N marker genes among the top drivers. Indeed, while in CT hCOs only the UL.Exc.N marker *PLXNA4* was found to drive differentiation trajectories, Het and Mos top driver genes included in addition *PLXNA4* (in Het only), *EPHA4*, *CUX2* and *SATB2* (the last two in Mos only, **Figure 2C**), suggesting a preferential and premature expression of the UL.Exc.N transcriptional program.

### Dysregulation of Notch and Wnt pathways in neural progenitors

To investigate whether altered cell type proportions in Het and Mos hCOs originate from progenitor structural defects, we analyzed the number and structure of neural rosettes - radially organized neuroepithelial structures that recapitulate the developing cortical ventricular zone. Quantification of neural rosettes in 1-month hCOs, identified by immunostaining for the progenitor marker SOX2, revealed significantly reduced rosette density in both Het (144 ± 72 rosettes/mm^2^) and Mos (118 ± 76 rosettes/mm^2^) compared to CT hCOs (350 ± 105 rosettes/mm^2^) (**Figure 3A)**. This reduction persisted in 3-month Mos hCOs (CT: 87 ± 53, Het: 48 ± 39, Mos: 28 ± 43 rosettes/mm^2^) and was rescued by rapamycin treatment (**Figure S3A and B**). Notably, rapamycin also increased rosette density in CT hCOs at 3 months (**Figure S3B**). Mos High hCOs showed a tendency towards lower density of rosettes compared to Mos Low hCOs at 1 month (**Figure S3C**), suggesting that the proportion of *DEPDC5*^-/-^ cells, and therefore the dosage of DEPDC5, plays a role in rosette formation and/or maintenance. Co-immunostaining for SOX2 and the tight-junction marker ZO1, which labels the ventricular lining, revealed no significant structural abnormalities in Het and Mos rosettes (**Figure 3B**). Furthermore, rosette ventricular size was comparable across all genotypes (**Figure S3D, E**). These data indicate that *DEPDC5* LoF and mTOR hyperactivity do not cause gross progenitor structural defects, and that the lower density of rosettes observed in Het and Mos may be a consequence rather than a cause of the accelerated differentiation revealed by scRNA-seq.

**Figure 3.**
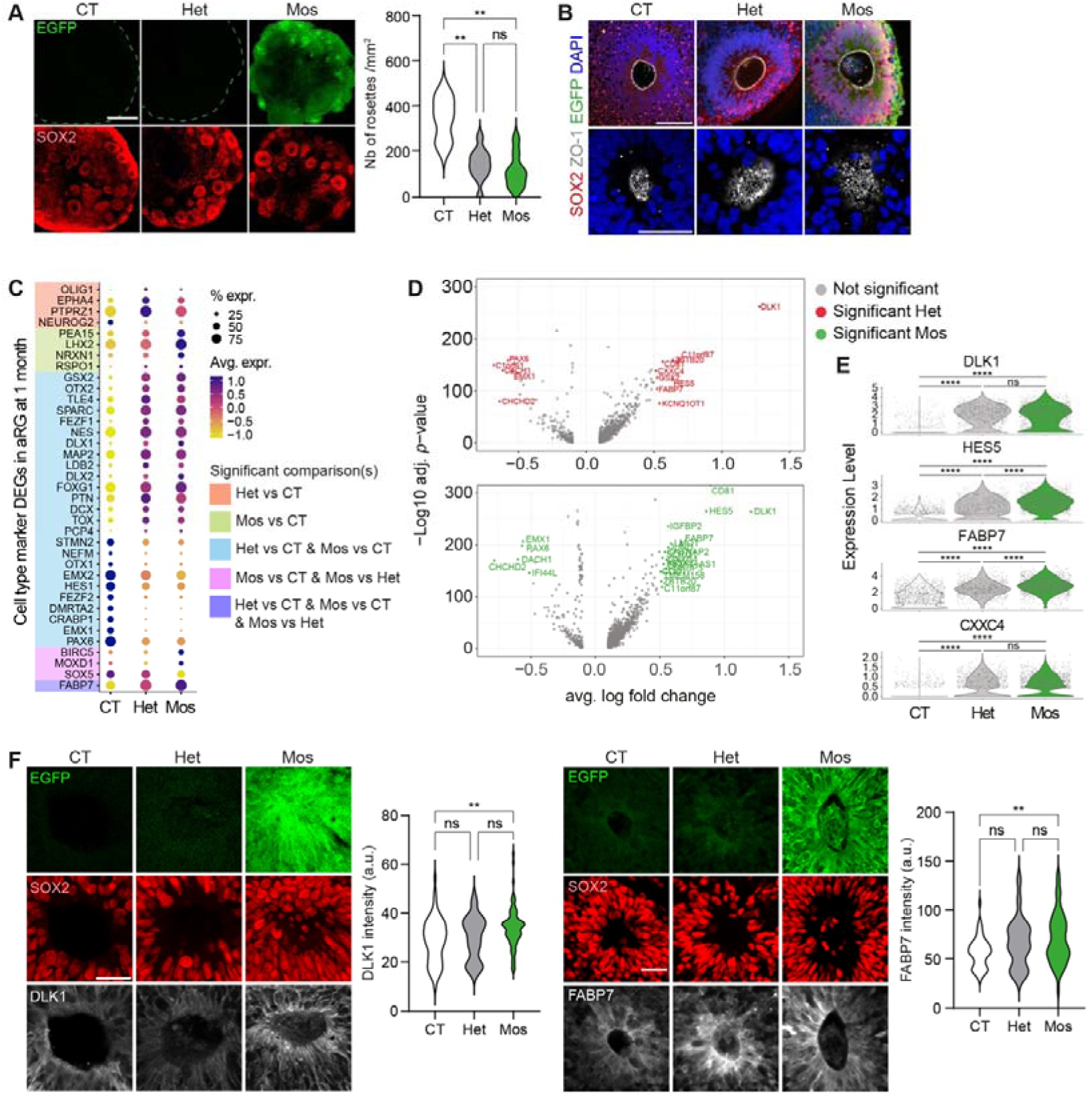
DEPDC5 deficiency alters developmental signaling pathways. (**A**) SOX2 and EGFP immunofluorescence in 1-month whole hCOs (confocal images). Scale bar: 500 µm. Rosette density quantified as number of rosettes/mm2. n = 3 CT, 10 Het, 8 Mos hCOs from 3 independent batches. One-way ANOVA with Dunnett’s correction. (**B**) Confocal z-stack analysis of neural rosettes immunostained for SOX2, ZO-1, and EGFP at 1 month. Upper: ZO-1 ring plane; Lower: ZO-1 surface plane. Scale bar, 50 µm. (**C**) Differential expression of cell type marker genes in apical radial glia (aRG) at 1 month. Comparisons between Het vs CT, Mos vs CT, and Mos vs Het. (**D**) Up-regulated and down-regulated genes (Wilcoxon Rank Sum test with Bonferroni correction) in Het and Mos 1-month aRGs compared to CT. Significance threshold was established as average log2FC > 0.5, adjusted p < 0.01. (based on Bonferroni correction using all genes in the dataset) < 0.01 (**E**) Expression levels of Notch (DLK1, HES5, FABP7) and Wnt (CXXC4) pathway genes in 1-month aRG across genotypes. (**F**) DLK1 and FABP7 immunofluorescence intensity quantified relative to SOX2+ rosette area (confocal images). Scale bar, 30 µm. ns, non-significant; \*\**p* < 0.01; \*\*\*\**p* < 0.0001.

To investigate transcriptional alterations in progenitor cells, we performed differential gene expression analysis comparing Het and Mos aRG with CT at 1 month. Differentially expressed genes (DEGs) included several known cell-type markers (**Figure 3C** and **Table S1**), suggesting that cell type specification changes may have emerged during the progenitor stage. Notably, both Het and Mos hCOs showed downregulation of aRG markers (*PAX6*, *EMX1* and *HES1)* as well as *FEZF2*, a key determinant of lower-layer subcortical projection neurons (**Figure 3C**). Mos-specific DEGs included progenitor markers such as *BIRC5*, *MOXD1*, *SOX5* and *FABP7* (**Figure 3C**).

We focused on DEGs with an absolute average log2 fold change (avg_Log2FC) > 0.5 and an adjusted *p*-value (p_val_adj) < 0.01 (**Figure 3D**). Among the top upregulated genes in both Het and Mos, we identified *DLK1*, *HES5*, *FABP7* and *CXXC4*, which are associated with Notch and Wnt signaling pathways (**Figure 3D and E**). Only *HES5* and *FABP7* were differentially expressed between Mos and Het aRG; however, the avg_Log2FC were lower than 0.5 (0.22 and 0.15, respectively; **Figure 3E**). Immunostaining against DLK1 and FABP7 confirmed their increased expression in 1-month Mos hCOs, but not Het (**Figure 3F**). Comparison of EGFP+ (DEPDC5^-/-^) and EGFP- (DEPDC5^+/-^) populations in Mos showed no significant differential expression of *DLK1*, *HES5*, *FABP7* or *CXXC4*. When comparing the two populations with CT and Het hCOs, the only difference came from *FABP7*, which was differentially expressed in the EGFP+ population but not in the EGFP-population compared to Het. The other three genes showed the same pattern in the two Mos populations in comparison to CT and Het (**Figure S3F** and **Table S1**). Notch and Wnt signaling pathways regulate cortical progenitor pool size, self-renewal versus differentiation balance, and neuronal laminar fate specification^29^. DLK1 promotes cell cycle exit in both mouse and human embryonic stem cell (ESC)-derived neural progenitors^30^. HES5 is a key effector of Notch signaling that regulates the transition timing of neurogenesis, with Hes5 overexpression in mouse cortical progenitors accelerating the transition from deep to superficial layer neurogenesis^31^. Our data indicate that *DEPDC5* loss in hCOs causes transcriptional upregulation of key Notch and Wnt signaling genes, which may underlie the premature differentiation of cortical progenitors and accelerated production of upper-layer excitatory neurons.

### *DEPDC5* mosaic two-hit hCOs exhibit altered synaptic gene expression and increased spontaneous neuronal spike activity

We then performed DEG analysis at more mature stages of development focusing on neuronal cells to identify dysregulated processes linked to FCDII pathophysiology. Given that our hCO model primarily recapitulates dorsal telencephalic development and exhibits limited interneuron specification, we restricted our analysis to early-born LL.Exc.Ns and late-born UL.Exc.N at timepoints corresponding to their peak abundance in CT hCOs, meaning 3 and 6 months, respectively (**Tables S2** and **S3**). LL.Exc.N exhibited a higher ratio of DEGs over expressed genes compared to UL.Exc.N in both Het and Mos organoids, suggesting increased vulnerability of early-born neurons to DEPDC5 loss (**Figure 4A**). Gene ontology (GO) enrichment analysis of DEGs for each excitatory neuron subtype revealed distinct enrichment patterns between genotypes. In Het LL.Exc.N, RNA processing and translation were predominantly affected (among the top 10 most enriched terms) while Mos LL.Exc.N showed enrichment of terms related to neuron projection and neuronal morphogenesis (**Figure 4B** and **Table S4**). RNA processing terms were also enriched when comparing Het to Mos LL.Exc.N (**Figure S4A** and **Table S4**), indicating a distinguishing feature between Het and Mos. Both Het and Mos UL.Exc.N also displayed altered expression of genes involved in neuron projection and neuronal morphogenesis compared to CT (**Figure 4B** and **Table S5**) but also when comparing Het and Mos (**Figure S4A** and **Table S5**). Genes associated with neuronal migration were specifically dysregulated in Mos UL.Exc.N (**Figure 4B**). These transcriptional alterations align with key histopathological features of FCDII, particularly defects in neuronal morphogenesis. The dysregulation of RNA metabolism and protein synthesis machinery in *DEPDC5* mosaic hCOs likely reflects downstream effects or compensatory mechanisms of activation of mTORC1 signaling.

**Figure 4.**
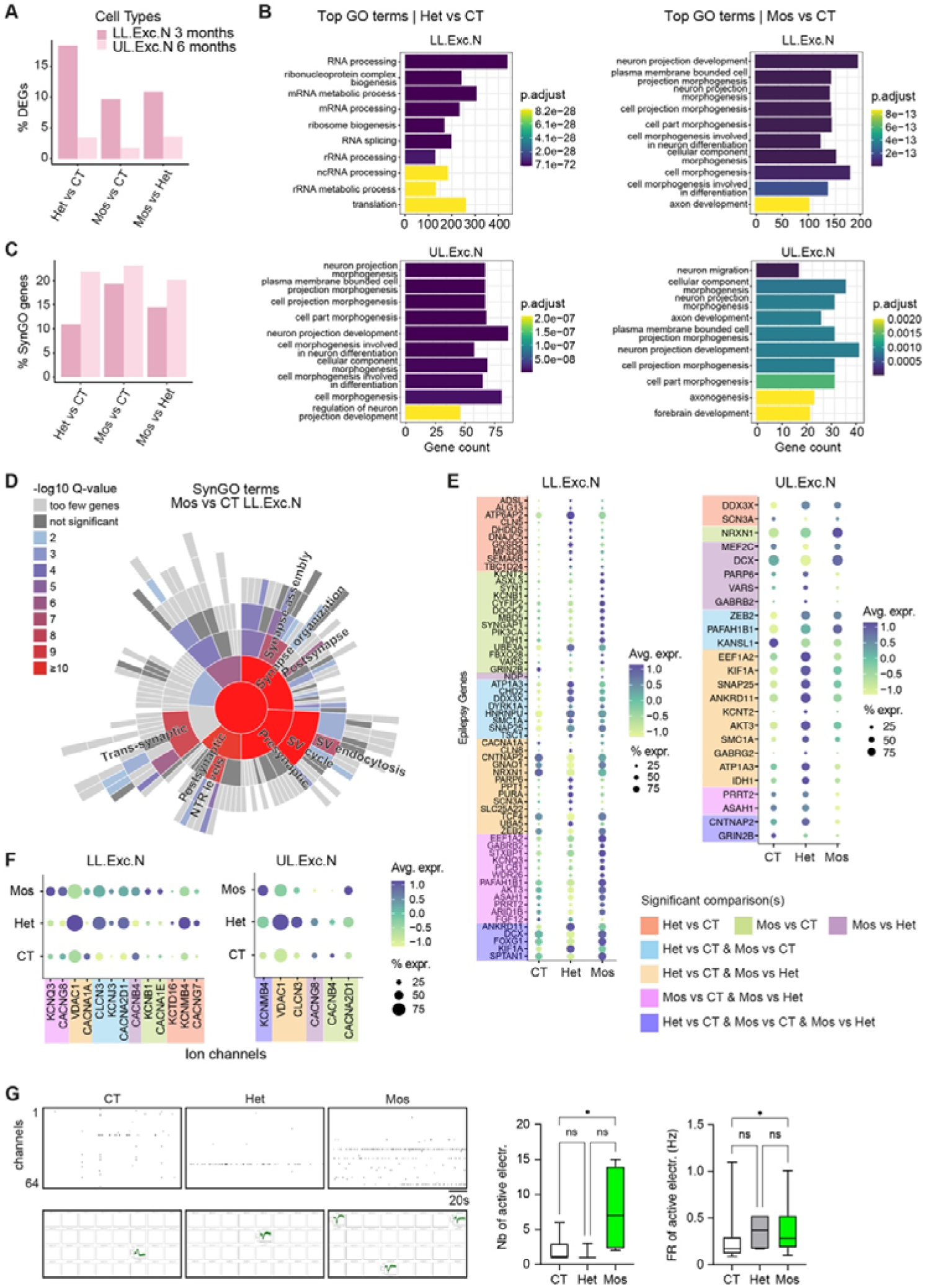
DEPDC5 deficiency alters synaptic gene expression and increases neuronal excitability. (**A**) Proportion of differentially expressed genes (DEGs) in lower-layer excitatory neurons (LL.Exc.Ns, 3 months) and upper-layer excitatory neurons (UL.Exc.Ns, 6 months). (**B**) Top 10 enriched GO biological process terms enriched in DEGs from LL.Exc.Ns and UL.Exc.Ns comparing Het and Mos hCOs to CT. (**C**) Proportion of synaptic genes (SynGO-annotated) among DEGs in LL.Exc.Ns (3 months) and UL.Exc.Ns (6 months) across genotype comparisons. (**D**) SynGO enrichment analysis of DEGs between Mos and CT LL.Exc.Ns at 3 months. Terms with log10 Q-value > 6 are shown. (**E**) Epilepsy-associated DEGs in Het and Mos LL.Exc.Ns and UL.Exc.Ns each versus CT, and Mos versus Het. (**F**) Ion-channel DEGs in Het and Mos LL.Exc.Ns and UL.Exc.Ns versus CT, and Mos versus Het. (**G**) Spontaneous electrical activity measured by multi-electrode array (MEA). Left: Representative raster plots showing spikes over 3 minutes and superimposed spikes from 32 electrodes. Right: Quantification of active electrodes (only organoids with at least one active electrode are listed here) and firing rate (FR). Box plots show median (middle line). Kruskal-Wallis test with Dunn’s correction. Active electrodes: n = 7 CT, 3 Het, 4 Mos organoids; Firing rate: n = 15 CT, 5 Het, 31 Mos electrodes from ≥3 independent batches. *p < 0.05.

Given that *DEPDC5* is a major epilepsy-causing gene, we next investigated potential molecular mechanisms underlying neuronal hyperexcitability and seizures in FCDII. We specifically examined differential expression of synaptic, epilepsy-related and ion channel genes. We interrogated the synaptic gene ontology (SynGO) database^32^, which catalogs 1,112 annotated synaptic genes (full SynGO analysis results are available in **Tables S6** and **S7**). Notably, in Mos hCOs, ∼20% of the DEGs in both UL.Exc.Ns and LL.Exc.Ns were SynGO-annotated genes. While Het UL.Exc.N showed comparable synaptic gene enrichment, Het LL.Exc.N exhibited only 10% match with SynGO-annotated genes (**Figure 4C**). Mos LL.Exc.Ns showed the highest Q-values in SynGO enrichment analysis compared to Mos UL.Exc.Ns, Het LL.Exc.Ns and Het UL.Exc.Ns (**Figures 4D** and **S4B**), with significant enrichment in genes controlling synapse organization, synaptic vesicle cycle and regulation of postsynaptic membrane neurotransmitter receptor levels (**Figure 4D**). We next interrogated DEGs for a comprehensive list of 143 established epilepsy-associated genes^33^. Of these genes, several matched with DEGs in both excitatory neuron subtypes, with LL.Exc.N showing the highest number of matching genes compared to UL.Exc.N. Nearly half of these genes were also differentially regulated between Het and Mos (**Figures 4E**). In addition, we found 13 and 6 DEGs matching ion channels in LL.Exc.Ns and UL.Exc.Ns, respectively (**Figure 4F**). Among these, three voltage-gated ion channel genes were previously linked to epilepsy: *KCNB1* (potassium voltage-gated channel subfamily B member 1) was specifically upregulated in Mos LL.Exc.N compared to CT, *CACNA1A* (calcium voltage-gated channel subunit alpha1 A) was downregulated in Het LL.Exc.N, and *KCNQ3* (potassium voltage-gated channel subfamily Q member 3) showed increased expression in Mos compared to both CT and Het (**Figure 4F**). We specifically interrogated ion channel genes expression in DEPDC5^+/-^ (EGFP-) and DEPDC5^-/-^ (EGFP+) within Mos cells compared to CT and Het hCOs, revealing a more severe dysregulation in EGFP+ cells (**Figure S4C**). Thus, Het and Mos excitatory neurons reveal dysregulation of genes and pathways potentially related to epilepsy insurgence in *DEPDC5*-related FCDII.

Several studies have shown that mature hCOs exhibit neuronal network activity^27,28,34^. To examine if the differential expression of epilepsy-associated genes affects the functional properties of neuronal networks in hCOs, we assessed spontaneous neuronal spike activity at 6 months using extracellular recording with a 64-electrode multi-electrode array (MEA). Our analysis focused on two key activity metrics: 1) the number of active electrodes and 2) the firing rate (FR). CT and Het hCOs displayed similarly low numbers (<5) of active electrodes (CT: 2.14 ± 0.71, n= 7; Het: 1.67 ± 0.67, n= 3). In contrast, Mos hCOs exhibited a significantly higher number of active electrodes compared to CT (7.75 ± 3.15, n= 4, p=0.049). Regarding the FR, CT and Het hCOs showed comparable values (0.26 ± 0.07 Hz, n=15 and 0.35 ± 0.08 Hz, n=5 respectively), and Mos were significantly higher than CT (0.39 ± 0.05; n=31, p=0.025) (**Figure 4G**).

In summary, although single-cell transcriptomic profiling revealed differential expression of synaptic, epilepsy-associated, and ion channel genes in both Het and Mos hCOs, only Mos hCOs exhibited hyperactive neuronal networks in MEA recording, suggesting that transcriptional changes in Het hCOs are insufficient to produce neuronal hyperexcitability, and biallelic *DEPDC5* inactivation is necessary for pathological network activity.

### DEPDC5 two-hit cells in mosaic hCOs exhibit metabolic and translational dysregulations

To examine cell-autonomous effects of biallelic DEPDC5 loss, we compared DEPDC5^-/-^ (EGFP+) and DEPDC5^+/-^ (EGFP-) cell populations within mosaic hCOs at both early and mature developmental stages. We first assessed whether EGFP+ cells participated in rosette formation to the same extent as EGFP-cells. Analysis of neural rosette composition in 1-month organoid sections revealed three distinct patterns: nearly entirely EGFP-rosettes, mosaic (EGFP- and EGFP+) rosettes, and nearly entirely EGFP+ rosettes (**Figure 5A**). Quantification of EGFP+ area within SOX2-rosettes demonstrated proportions consistent with the expected mosaicism rates of ∼50% and ∼20% in High Mos and Low Mos hCOs, respectively (**Figure S5A**). These findings indicate that DEPDC5^-/-^ cells integrate into rosettes without bias, suggesting that biallelic *DEPDC5* loss does not significantly alter early neuroepithelial organization.

**Figure 5.**
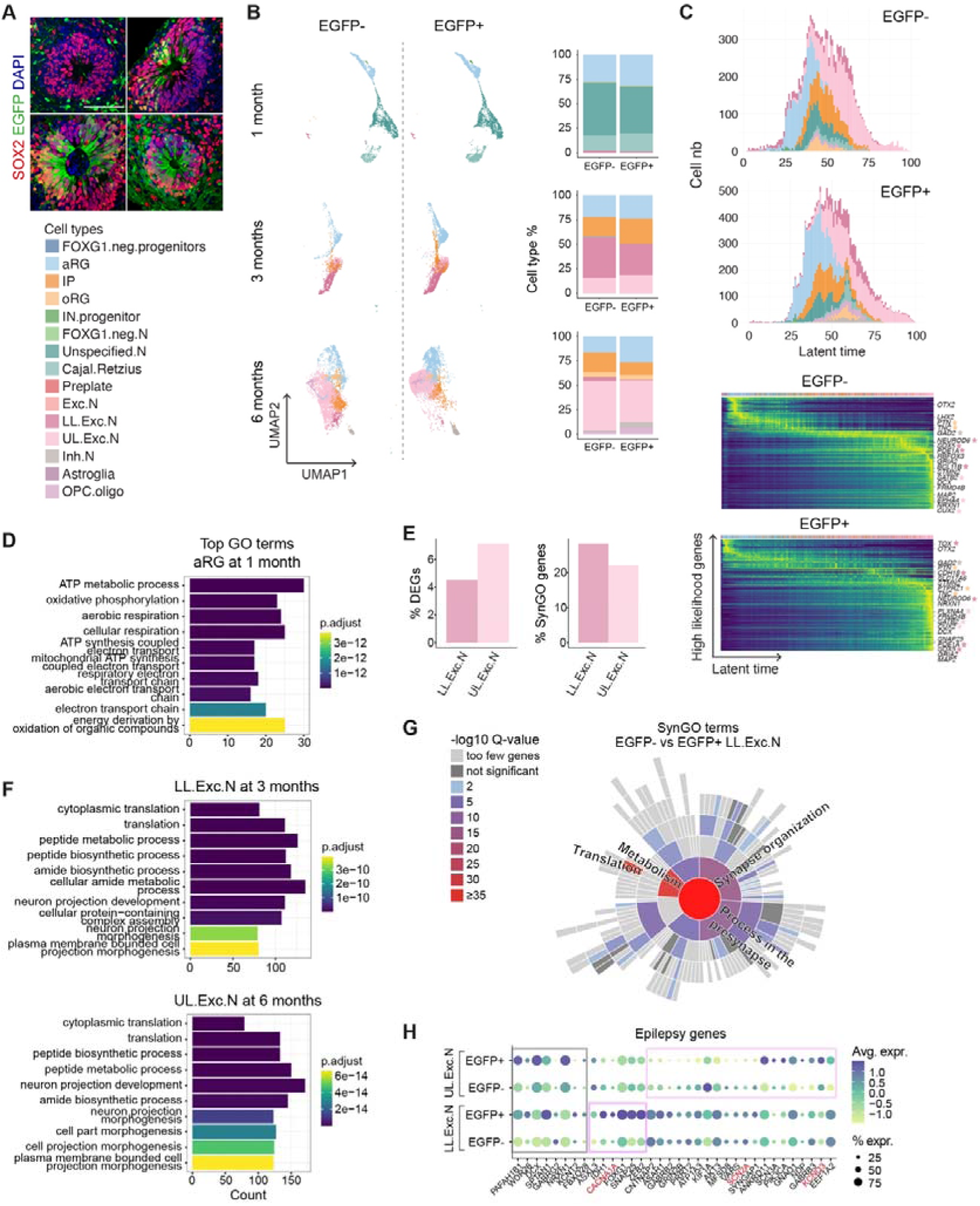
Comparison of *DEPDC5*^+/-^ and *DEPDC5*^-/-^ cell populations in mosaic cortical organoids. (**A**) SOX2 and EGFP immunofluorescence in 1-month Mos hCOs, showing three rosette types: entirely *DEPDC5*^+/-^ EGFP-rosettes (top left), mosaic (EGFP- and *DEPDC5*^-/-^ EGFP+) rosettes (top right), and nearly entirely EGFP+ rosettes (bottom left and right). (**B**) Single-cell transcriptomic analysis of EGFP+ and EGFP-populations in Mos hCOs. UMAP visualizations showing conserved cell clusters between EGFP+ and EGFP-populations. Right: Cell type composition at each timepoint. EGFP+ defined as cells with EGFP counts > 0. (**C**) RNA velocity analysis comparing EGFP+ and EGFP-populations at all timepoints. Top: Cell type evolution along latent time. Bottom: Heatmaps representing gene expression dynamics resolved along latent time show the cascade of transcription for the top 300 likelihood-ranked genes. Colored stars indicate cell type-specific markers. (**D**) Top 10 most-significant GO biological process terms enriched in DEGs between EGFP+ and EGFP-aRG at 1 month. (**E**) Differential gene expression analysis between EGFP+ and EGFP-populations. Left: Proportion of DEGs in LL.Exc.Ns (3 months) and UL.Exc.Ns (6 months). Right: Proportion of synaptic genes (SynGO-annotated) among DEGs. (**F**) Top 10 most-significant GO biological process terms enriched in DEGs between EGFP+ and EGFP-LL.Exc.Ns. (log10 Q-value > 10 shown). (**G**) SynGO enrichment analysis of DEGs between EGFP+ and EGFP-LL.Exc.Ns (log10 Q-value > 10 shown). (**H**) Epilepsy-associated genes DEGs between EGFP+ and EGFP-LL.Exc.Ns and UL.Exc.Ns. Ion channel genes indicated in red.

To assess whether developmental alterations were cell-autonomous, we analyzed EGFP+ and EGFP-cell populations within Mos hCO using scRNA-seq data. Cell type proportions showed only minor differences between EGFP+ and EGFP- populations across all timepoints (1, 3 and 6 months; **Figure 5B**). scVelo trajectory analysis confirmed similar cell type differentiation dynamics between both populations (**Figure 5C**). These findings indicate that the altered cell type compositions observed in Mos hCOs (**Figure 2A**) likely arise through non-cell-autonomous mechanisms.

To identify potential transcriptomic differences between DEPDC5^+/-^ and DEPDC5^-/-^ cell populations, we conducted DGE and GO analyses across key developmental stages and cell types (**Tables S1 to S7**), focusing as above on aRG at 1 month, LL.Exc.Ns at 3 months, and UL.Exc.Ns at 6 months. aRG showed alterations of metabolic pathways, particularly ATP metabolism, oxidative phosphorylation, and cellular respiration (**Figure 5D**). UL.Exc.Ns showed a higher percentage of DEGs compared to LL.Exc.Ns (**Figure 5E**), with top GO terms such as translation, metabolic and neuron projection development in both excitatory neuron subtypes (**Figure 5F**). A substantial proportion of DEGs in both LL.Exc.N (28%) and UL.Exc.N (22%) mapped to SynGO (**Figure 5E**). Among these SynGO-annotated genes, 45% in LL.Exc.N and 19% in UL.Exc.N overlapped with those identified in the Mos versus CT hCO comparison. Consistent with GO enrichment analysis, SynGO enrichment analysis revealed predominant dysregulation of metabolic processes, synaptic translation as well as synapse organization in both LL.Exc.N and UL.Exc.N (**Figure 5G** and **Figure S5B**). Among the DEGs in the two excitatory neuron subtypes, we identified several epilepsy-associated genes, including LL.Exc.Ns (71%) and UL.Exc.Ns (21%) which were also found in the Mos versus CT comparison (**Figure 5H**). Notable dysregulated ion channel genes, previously linked to epilepsy, included *KCNQ3* (upregulated in DEPDC5^-/-^ UL.Exc.Ns), *CACNA1A* (upregulated in DEPDC5^-/-^ in LL.Exc.Ns), and SCN2A (downregulated in DEPDC5^-/-^ in UL.Exc.Ns). These cell-autonomous transcriptional changes affecting *DEPDC5*^-/-^ compared to *DEPDC5*^+/-^ excitatory neurons may exacerbate the Mos phenotype and contribute to the epileptogenic nature of FCDII lesions harboring biallelic DEPDC5 loss.

## Discussion

In this study, we leveraged human cortical organoids (hCOs) to model both heterozygous (Het) and biallelic loss-of-function *DEPDC5* mutations in a mosaic context (Mos), providing insights into the role of DEPDC5 and its regulation of mTOR activity in neurodevelopment. Through longitudinal data across three developmental timepoints, we identified early neurodevelopmental phenotypes in both Het and Mos hCOs including altered neural rosette density and premature generation of upper-layer neurons, which were associated with dysregulation of Notch and Wnt signaling pathways. Furthermore, our findings, provide experimental support for Knudson’s two-hit model in FCDII surgical tissues where a second-hit somatic mutation leading to biallelic *DEPDC5* inactivation is required for the FCDII phenotype including mTOR-hyperactive dysmorphic-like neurons and spontaneous neuronal network hyperactivity, which were absent in Het hCO^24^. Notably, *DEPDC5* mosaic organoids did not develop balloon cells, consistent with clinical observations that *DEPDC5* mutations predominantly cause FCDIIa, characterized by dysmorphic neurons without balloon cells^9,21^. The necessity of a second hit for the development of pS6-positive dysmorphic neurons was also observed in another mTORopathy cortical organoid model of Tuberous Sclerosis Complex (TSC) with *TSC1/TSC2* mutations^35^.

Patients with germline heterozygous *DEPDC5* mutations and nonlesional epilepsy rarely undergo brain surgery, limiting opportunities for histopathological and genetic analyses of brain tissue. Thus, it remains unclear whether single-hit heterozygous *DEPDC5* loss-of-function alone is sufficient to generate seizures. Our electrophysiological findings align with mouse models, where homozygous deletion causes spontaneous seizures while heterozygous deletion reduces seizure threshold^36–40^. Consistent with this observation, the increased firing rate in both Het and Mos hCOs suggests that loss of heterozygosity can confer seizure susceptibility^35^. We identified transcriptional alterations in excitatory neuron subtypes shared between Het and Mos, as well as specific to each genotype including altered expression of synaptic and ion channel genes, with KCNQ3 specifically upregulated in double-hit cells. Although our data do not allow us to clearly identify which gene(s) may be hold responsible for hyperactivity, we point towards several epilepsy-associated and ion channel genes that are more severely affected in Mos.

Our mosaic organoid model enabled us to distinguish cell-autonomous from non-cell-autonomous effects of biallelic *DEPDC5* loss. Comparison of DEPDC5^+/-^ and DEPDC5^-/-^ cells within Mos hCOs suggests that altered development is predominantly driven by non-cell-autonomous mechanisms, suggesting a scenario where global mTOR activity levels influence timing of fate specification in the developing cerebral cortex. On the other hand, cell-autonomous effects mostly pointed towards alterations in metabolism (including cell respiration processes) and translation, which likely drive the formation of cytomegalic dysmorphic neurons, although a clear mechanistic link needs providing.

Our study has some limitations. First, our findings are based on a single *DEPDC5* patient-derived line, necessitating validation in additional *DEPDC5* two-hit iPSC lines to establish phenotypic reproducibility. Second, the use of homozygous DEPDC5 knockout hiPSCs may not accurately model the temporal dynamics of somatic mutation acquisition in patients, where variants likely arise during later developmental stages. This early depletion could trigger compensatory mechanisms that potentially attenuate the observed phenotypes. Third, we were unable to generate an isogenic control. However, some of our key findings rely on comparisons between DEPDC5^+/-^ and DEPDC5^-/-^ cells within the same mosaic organoids, mitigating the lack of an external isogenic control. Future studies employing inducible *DEPDC5* knockout systems and additional patient-derived lines will be important to address these limitations.

Despite these limitations, our study sheds light on the molecular mechanisms underlying DEPDC5-related FCDII, demonstrating the utility of mosaic hCOs in modeling focal cortical malformations. We provide here the bases for future studies aiming at identifying potential therapeutic targets among the identified dysregulated pathways. These advances will pave the way for more personalized approaches in understanding and treating DEPDC5-related cortical malformations and epilepsies.

## Acknowledgements

We thank the ICM core facilities: for imaging (ICM.Quant), for histology (Histomics), for sequencing (iGenSeq), for bioinformatic analysis (DAC), for DNA and cell bank, for FANS-sorting (ICV-3C) and for hIPSCs (iPS-CELIS). We thank Manon Quiquand and Eric Noé for technical assistance, and Sara Baldassari and Ann-Sofie de Meulemeester for helpful feedback on the manuscript. We thank the patients and their families

## Author contributions

M.M., T.R. and S.Bau. conceived the study. M.M. and T.R. generated human-induced pluripotent stem-cell lines and performed CRISPR-Cas9 genome editing. M.M. led and performed most of the experimental work, including generation and maintenance of human cortical organoids and hiPSCs, immunostaining, microscopy and image analysis, western-blotting and related quantifications, and single-cell RNA-sequencing sample preparation. T.R., K.G. and M.D. assisted with and performed generation and maintenance of human cortical organoids, sectioning and immunostaining, microscopy and image analysis, and western blotting. S.B. helped with organoid culture and maintenance. S.B. led and performed scRNA-seq bioinformatic analyses. C.R. developed the scRNA-seq bioinformatics core pipeline and performed the RNA velocity analyses. C.D. performed multielectrode array electrophysiology experiments and data analysis. F.B. recruited the patients. M.M. and S.B. generated manuscript figures. M.M., S.B. and S. Bau interpreted the data and wrote the manuscript. S.Bau. directed the research.

## SUPPLEMENTARY FIGURES

**Figure S1.**
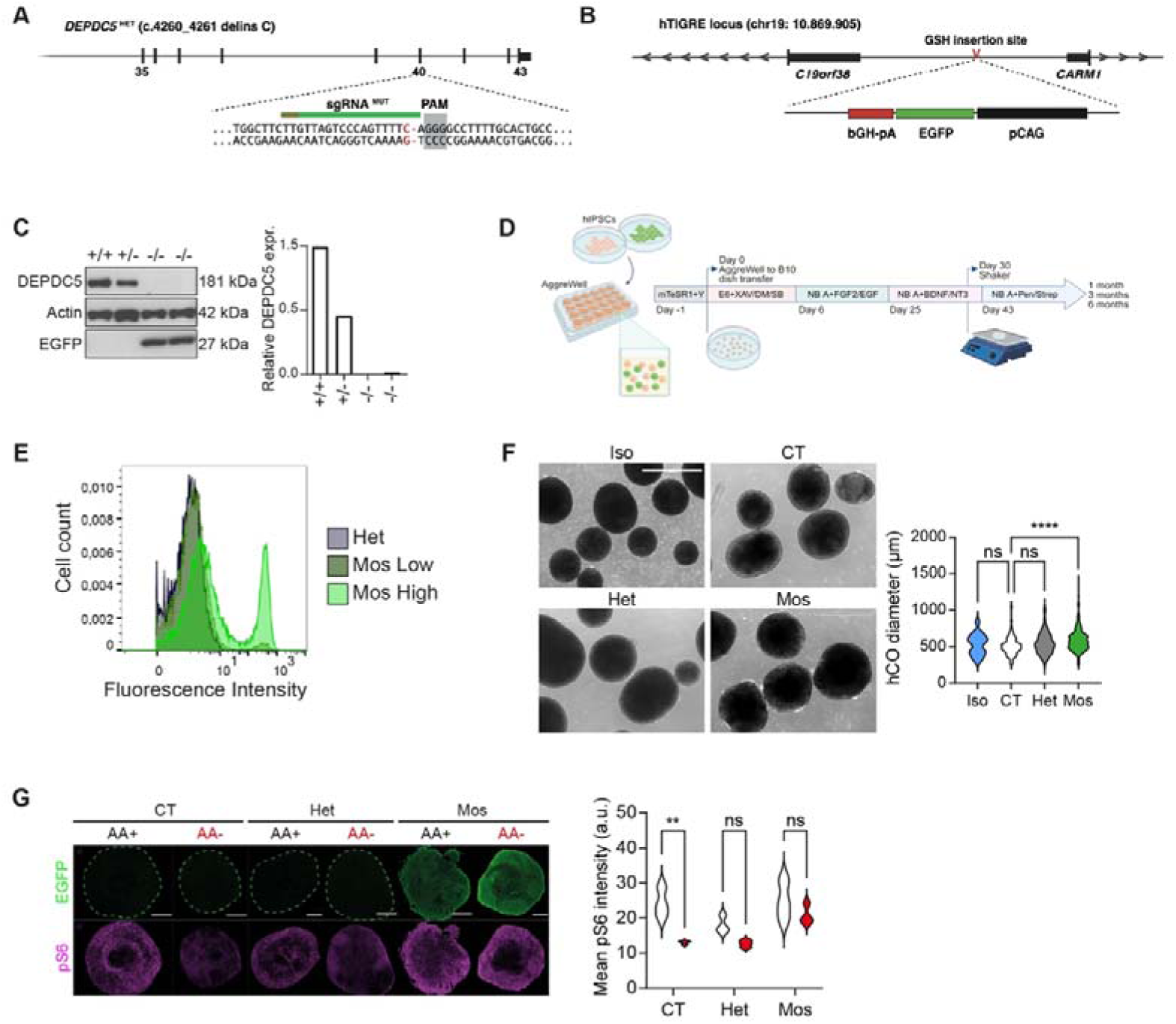
Generation and validation of DEPDC5-/- hiPSC lines and cortical organoids. **(A)** CRISPR-Cas9 targeting strategy for DEPDC5 exon 40 to introduce p.L1420Ffs*154 mutation in DEPDC5+/- hiPSCs. PAM: protospacer adjacent motif. (**B**) Strategy for EGFP insertion into hTIGRE locus to generate EGFP-tagged DEPDC5-/- line. (**C**) Western blot of DEPDC5, EGFP, and actin in *DEPDC5^+/+^*, *DEPDC5^+/-^*, *DEPDC5^-/-^* (clone 1), and *DEPDC5^-/-^* (clone 2) hiPSC lines. Protein levels normalized to actin and control levels. (**D**) Protocol for generating mosaic organoids (Mos hCO). EGFP+ and EGFP-hiPSCs mixed at defined ratios in AggreWell plates. Timeline shows media composition and cytokine treatments, as described in^26^. (**E**) Flow cytometric analysis of EGFP expression in 1-month Het and Mos hCOs. (**F**) Day 14 organoid morphology and size quantification (bright-field images). Scale bar, 1000 µm. n = 16 Iso, 94 CT, 119 Het, 316 Mos High hCOs (from one batch). Kruskal-Wallis test with Dunn’s correction. (**G**) GFP and pS6 immunofluorescence in 6-month hCO sections under basal (AA+) and amino acid-deprived (AA-) conditions. pS6 intensity normalized to section area. Scale bar, 500 µm. n = 3 organoids/condition, one section/organoid. Two-way ANOVA with Šidák’s correction. ****, p < 0.001.

**Figure S2.**
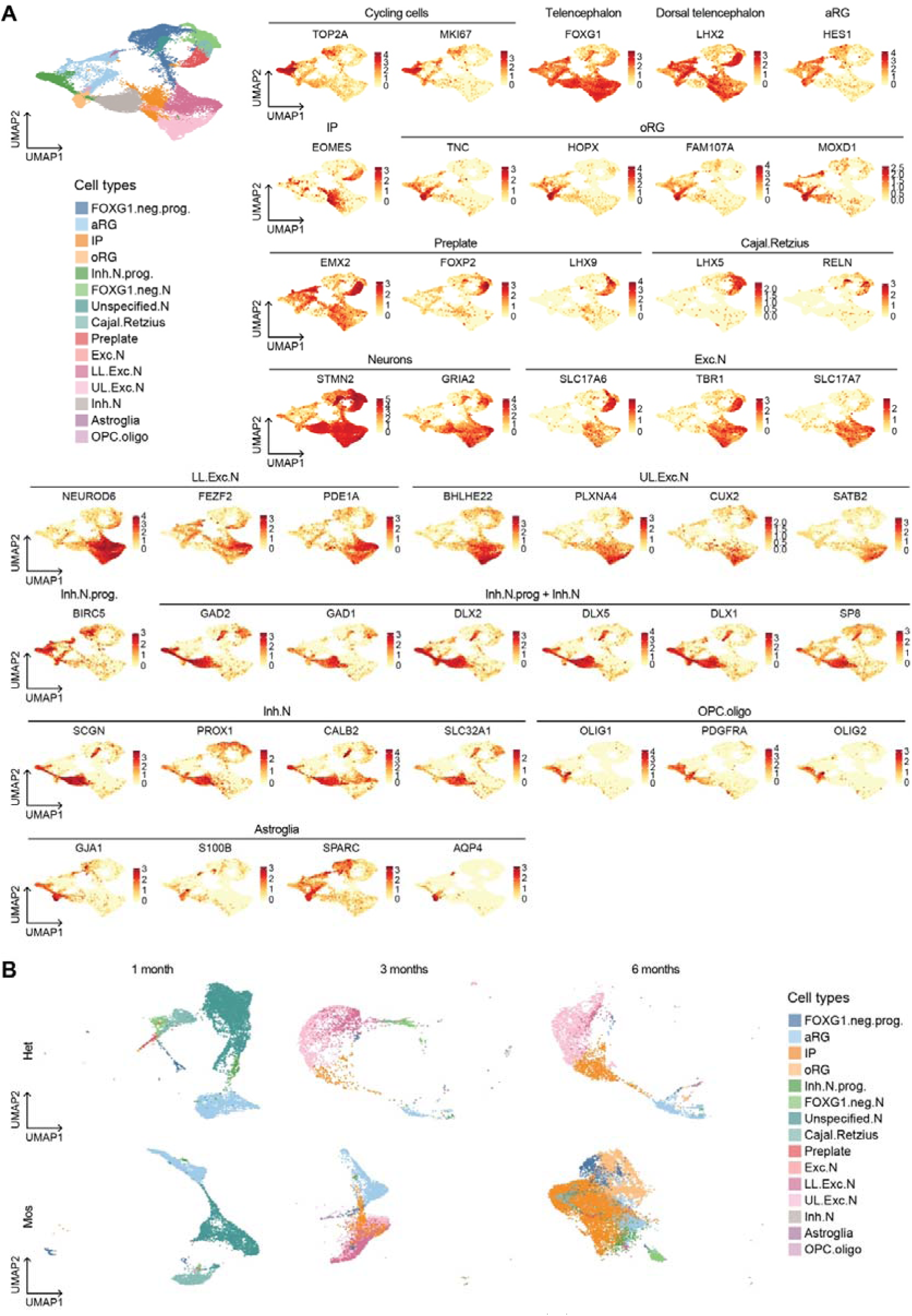
Cell type annotation from scRNA-seq. (**A**) Expression of canonical cell type markers in CT hCOs. UMAP and feature plots represent the integration of scRNA-seq data obtained at 1, 3 and 6 months. (**B**) Cell types obtained by label transfer from CT in Het and Mos hCOs.

**Figure S3.**
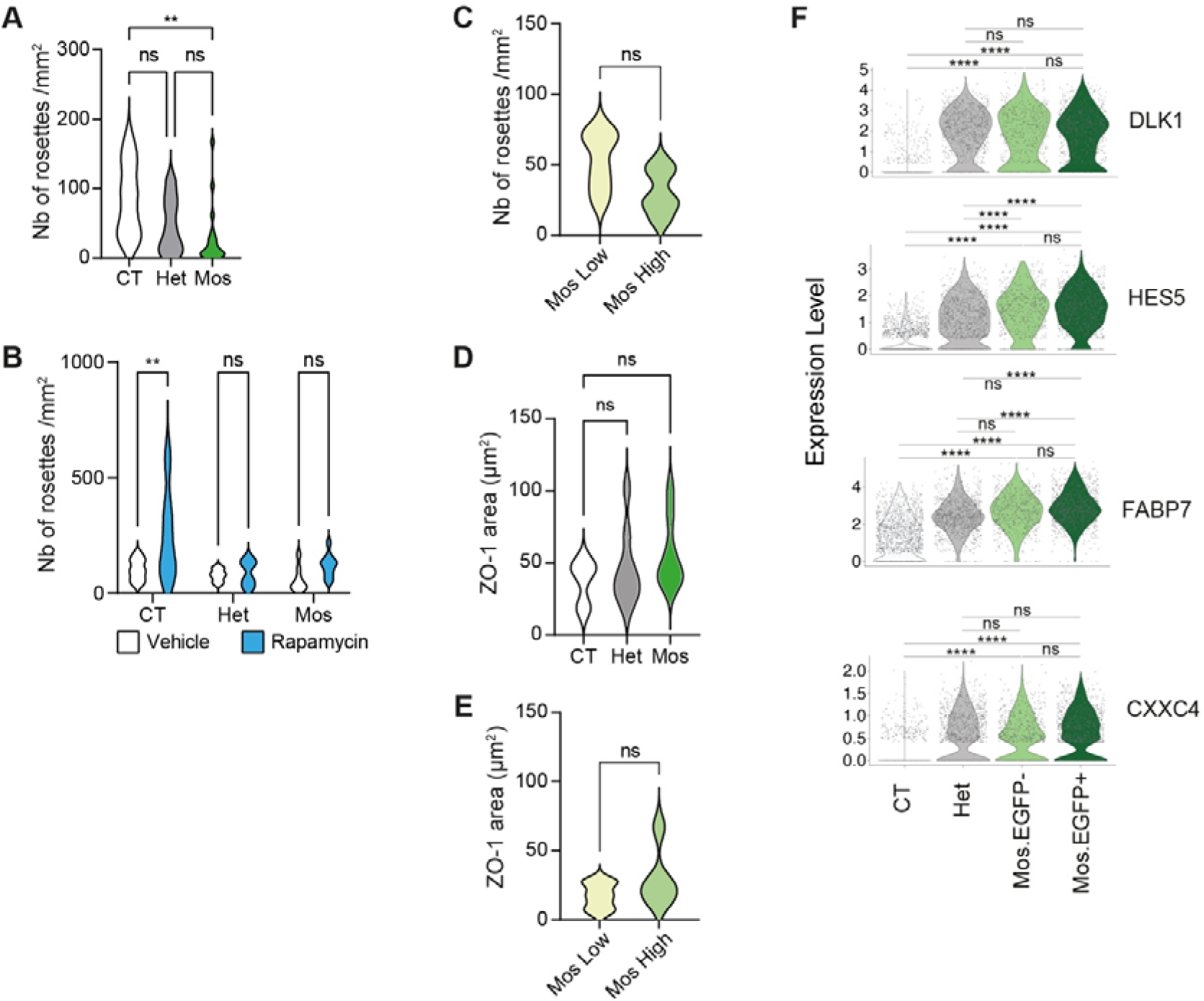
Characterization of neural rosette formation across DEPDC5 genotypes. (**A**) Rosette density at 3 months normalized to section area. n = 10 CT, 15 Het, 18 Mos organoids from 3 independent batches, one section/hCO. Kruskal-Wallis test with Dunn’s correction. **(B)** Impact of rapamycin treatment on rosette density at 3 months. n = 3-5 organoids/genotype/treatment (from one batch.) Two-way ANOVA with Šidák’s correction. **(C)** Comparison of rosette density between Mos Low and Mos High organoids at 1 month. N = 6 Mos Low, 5 Mos High organoids from 2 independent batches. Mann-Whitney test. (**D**) ZO-1 area quantification at 1 month. n = 3 CT, 9 Het, 8 Mos hCOs from 3 independent batches. Kruskal-Wallis test with Dunn’s correction. (**E**) ZO-1 area comparison between Mos Low and Mos High hCO at 1 month. n = 6 Mos Low, 5 Mos High organoids from 2 independent batches. Mann-Whitney test. ns, non-significant; **, *p* < 0.01. (**F**) Expression of Notch (DLK1, HES5, FABP7) and Wnt (CXXC4) pathway genes in 1-month aRG comparing Mos EGFP- and EGFP+.

**Figure S4.**
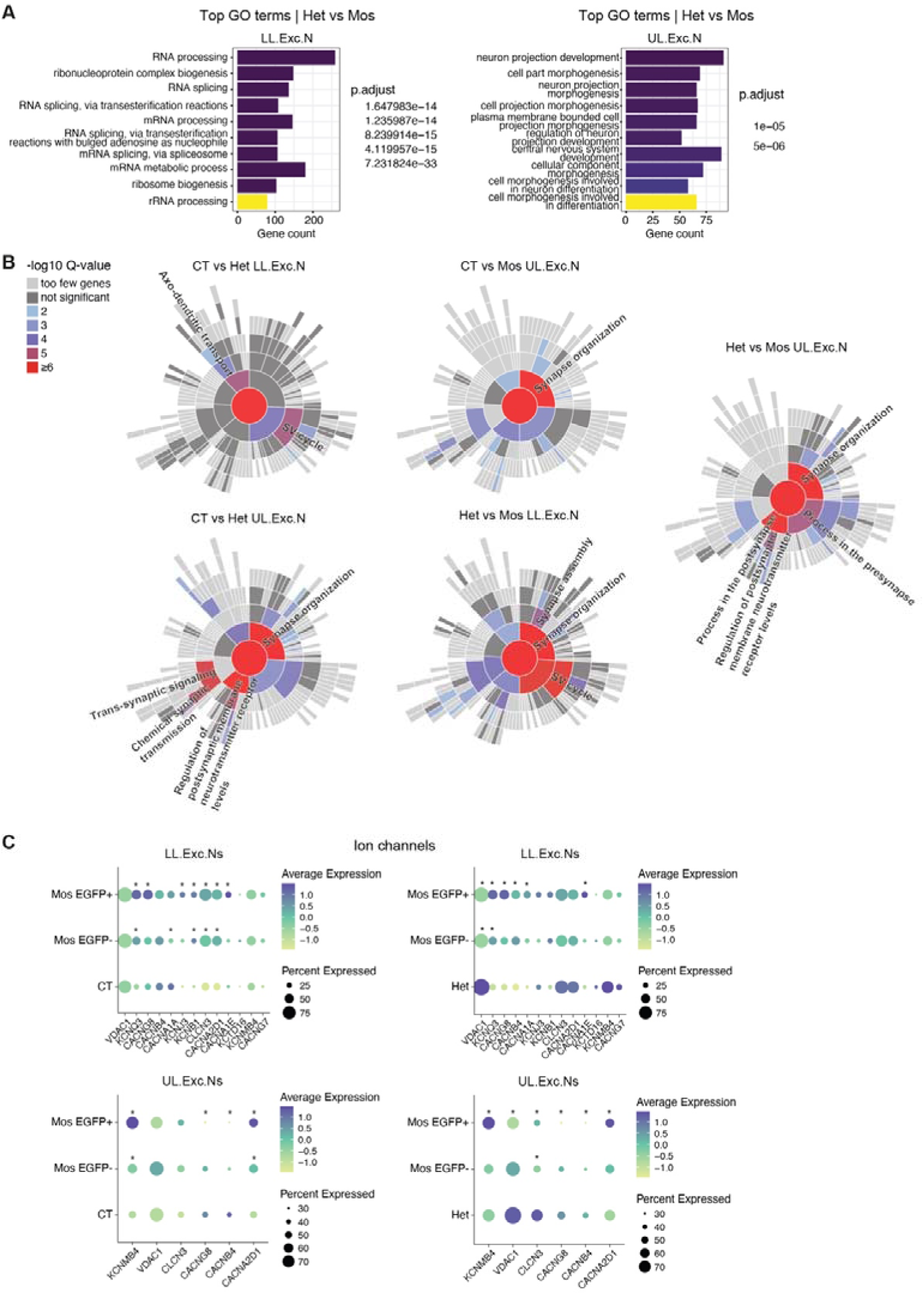
Gene ontology and SynGO enrichment analyses. (**A**) Top 10 most significant GO biological process terms enriched in DEGs between Mos and Het in lower-layer (LL.Exc.Ns) and upper-layer (UL.Exc.Ns) excitatory neurons. (**B**) SynGO enriched biological process terms. (**C**) Ion channel gene expression profiles in EGFP- and EGFP+ populations from Mos hCOs compared to CT and Het, analyzed in LL.Exc.Ns at 3 months and UL.Exc.Ns at 6 months.

**Figure S5.**
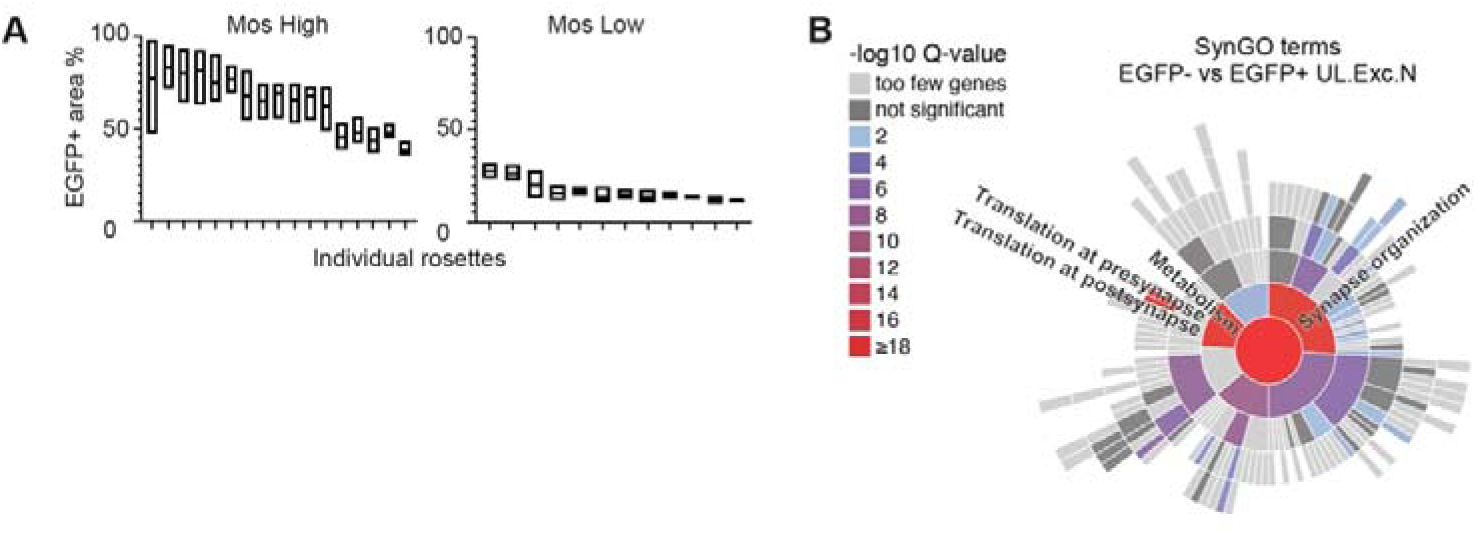
Comparison of *DEPDC5*^+/-^ and *DEPDC5*^-/-^ cells in Mos hCO. (**A**) Quantification of EGFP+ area within individual rosettes at 1 month. n = 17, high Mos, n= 12, low Mos hCOs. (**B**) SynGO biological process terms enriched for DEGs between EGFP+ and EGFP-UL.Exc.Ns in Mos hCOs.

## Supplementary Tables

**<u>Table S1.</u>** Full list of differentially expressed genes in aRG at 1 month.

**<u>Table S2.</u>** Full list of differentially expressed genes in LL.Exc.Ns at 3 months.

**<u>Table S3.</u>** Full list of differentially expressed genes in UL.Exc.Ns at 6 months.

**<u>Table S4.</u>** Gene ontology enrichment analysis performed on differentially expressed genes in LL.Exc.N at 3 months.

**<u>Table S5.</u>** Gene ontology enrichment analysis performed on differentially expressed genes in UL.Exc.N at 6 months.

**<u>Table S6.</u>** Full results of SynGO analysis for LL.Exc.N at 3 months.

**<u>Table S7.</u>** Full results of SynGO analysis for UL.Exc.N at 6 months.

## References

1. Blumcke, I., Spreafico, R., Haaker, G., Coras, R., Kobow, K., Bien, C.G., Pfafflin, M., Elger, C., Widman, G., Schramm, J., et al. (2017). Histopathological Findings in Brain Tissue Obtained during Epilepsy Surgery. The New England journal of medicine 377, 1648–1656. 10.1056/NEJMoa1703784.

2. Najm, I., Lal, D., Alonso Vanegas, M., Cendes, F., Lopes-Cendes, I., Palmini, A., Paglioli, E., Sarnat, H.B., Walsh, C.A., Wiebe, S., et al. (2022). The ILAE consensus classification of focal cortical dysplasia: An update proposed by an ad hoc task force of the ILAE diagnostic methods commission. Epilepsia 63, 1899–1919. 10.1111/epi.17301.

3. Macdonald-Laurs, E., Warren, A.E.L., Lee, W.S., Yang, J.Y., MacGregor, D., Lockhart, P.J., Leventer, R.J., Neal, A., and Harvey, A.S. (2023). Intrinsic and secondary epileptogenicity in focal cortical dysplasia type II. Epilepsia 64, 348–363. 10.1111/epi.17495.

4. Corrigan, R.R., Mashburn-Warren, L.M., Yoon, H., and Bedrosian, T.A. (2024). Somatic Mosaicism in Brain Disorders. Annu Rev Pathol. 10.1146/annurev-pathmechdis-111523-023528.

5. Bizzotto, S., and Walsh, C.A. (2022). Genetic mosaicism in the human brain: from lineage tracing to neuropsychiatric disorders. Nature reviews. Neuroscience 23, 275–286. 10.1038/s41583-022-00572-x.

6. Blumcke, I., Budday, S., Poduri, A., Lal, D., Kobow, K., and Baulac, S. (2021). Neocortical development and epilepsy: insights from focal cortical dysplasia and brain tumours. Lancet Neurol 20, 943–955. 10.1016/S1474-4422(21)00265-9.

7. Lamparello, P., Baybis, M., Pollard, J., Hol, E.M., Eisenstat, D.D., Aronica, E., and Crino, P.B. (2007). Developmental lineage of cell types in cortical dysplasia with balloon cells. Brain 130, 2267–2276. 10.1093/brain/awm175.

8. Baldassari S, Klingler E, Gomez Teijeiro L, Doladilhe M, Raoux C, Roig Puiggro S, Bizzotto S, Sami L, Ribierre T, Aronica E, Biassette H, Chipaux M, Jabaudon D and Baulac S (2025). Single-cell genotyping and transcriptomic profiling in focal cortical dysplasia. Nat neuroscience 2025 May;28(5):964–972. 10.1038/s41593-025-01936-z.

9. Gerasimenko, A., Baldassari, S., and Baulac, S. (2023). mTOR pathway: Insights into an established pathway for brain mosaicism in epilepsy. Neurobiol Dis 182, 106144. 10.1016/j.nbd.2023.106144.

10. D’Gama, A.M., Woodworth, M.B., Hossain, A.A., Bizzotto, S., Hatem, N.E., LaCoursiere, C.M., Najm, I., Ying, Z., Yang, E., Barkovich, A.J., et al. (2017). Somatic Mutations Activating the mTOR Pathway in Dorsal Telencephalic Progenitors Cause a Continuum of Cortical Dysplasias. Cell reports 21, 3754–3766. 10.1016/j.celrep.2017.11.106.

11. Baldassari, S., Ribierre, T., Marsan, E., Adle-Biassette, H., Ferrand-Sorbets, S., Bulteau, C., Dorison, N., Fohlen, M., Polivka, M., Weckhuysen, S., et al. (2019). Dissecting the genetic basis of focal cortical dysplasia: a large cohort study. Acta neuropathologica 138, 885–900. 10.1007/s00401-019-02061-5.

12. Sim, N.S., Ko, A., Kim, W.K., Kim, S.H., Kim, J.S., Shim, K.W., Aronica, E., Mijnsbergen, C., Spliet, W.G.M., Koh, H.Y., et al. (2019). Precise detection of low-level somatic mutation in resected epilepsy brain tissue. Acta neuropathologica 138, 901–912. 10.1007/s00401-019-02052-6.

13. Chung, C., Yang, X., Bae, T., Vong, K.I., Mittal, S., Donkels, C., Westley Phillips, H., Li, Z., Marsh, A.P.L., Breuss, M.W., et al. (2023). Comprehensive multi-omic profiling of somatic mutations in malformations of cortical development. Nature genetics 55, 209–220. 10.1038/s41588-022-01276-9.

14. Pirozzi, F., Berkseth, M., Shear, R., Gonzalez, L., Timms, A.E., Sulc, J., Pao, E., Oyama, N., Forzano, F., Conti, V., et al. (2022). Profiling PI3K-AKT-MTOR variants in focal brain malformations reveals new insights for diagnostic care. Brain 145, 925–938. 10.1093/brain/awab376.

15. Panwar, V., Singh, A., Bhatt, M., Tonk, R.K., Azizov, S., Raza, A.S., Sengupta, S., Kumar, D., and Garg, M. (2023). Multifaceted role of mTOR (mammalian target of rapamycin) signaling pathway in human health and disease. Signal Transduct Target Ther 8, 375. 10.1038/s41392-023-01608-z.

16. Baldassari, S., Picard, F., Verbeek, N.E., van Kempen, M., Brilstra, E.H., Lesca, G., Conti, V., Guerrini, R., Bisulli, F., Licchetta, L., et al. (2018). The landscape of epilepsy-related GATOR1 variants. Genetics in medicine: official journal of the American College of Medical Genetics. 10.1038/s41436-018-0060-2.

17. Samanta, D. (2022). DEPDC5-related epilepsy: A comprehensive review. Epilepsy & behavior: E&B 130, 108678. 10.1016/j.yebeh.2022.108678.

18. Ochoa-Urrea, M., Butler, E.A., Brunger, T., Bosselmann, C., Najm, I., Lhatoo, S.D., and Lal, D. (2024). Insights into *DEPDC5*-Related Epilepsy from 586 people: Variant Penetrance, Phenotypic Spectrum, and Treatment Outcomes. medRxiv, 2024.2012.2025.24319647. 10.1101/2024.12.25.24319647.

19. Epi, C. (2024). Exome sequencing of 20,979 individuals with epilepsy reveals shared and distinct ultra-rare genetic risk across disorder subtypes. Nature neuroscience 27, 1864–1879. 10.1038/s41593-024-01747-8.

20. Baldassari, S., Picard, F., Verbeek, N.E., van Kempen, M., Brilstra, E.H., Lesca, G., Conti, V., Guerrini, R., Bisulli, F., Licchetta, L., et al. (2019). The landscape of epilepsy-related GATOR1 variants. Genet Med 21, 398–408. 10.1038/s41436-018-0060-2.

21. Honke, J., Hoffmann, L., Coras, R., Kobow, K., Leu, C., Pieper, T., Hartlieb, T., Bien, C.G., Woermann, F., Cloppenborg, T., et al. (2023). Deep histopathology genotype-phenotype analysis of focal cortical dysplasia type II differentiates between the GATOR1-altered autophagocytic subtype IIa and MTOR-altered migration deficient subtype IIb. Acta Neuropathol Commun 11, 179. 10.1186/s40478-023-01675-x.

22. Baulac, S., Ishida, S., Marsan, E., Miquel, C., Biraben, A., Dang Khoa, N., Nordli, D., Cossette, P., Sylvie, N., Lambrecq, V., et al. (2015). Familial Focal Epilepsy with Focal Cortical Dysplasia Due to DEPDC5 Mutations. Annals of Neurology 77, 675–683. 10.1002/ana.24368.

23. Mirzaa, G.M., Campbell, C.D., Solovieff, N., Goold, C., Jansen, L.A., Menon, S., Timms, A.E., Conti, V., Biag, J.D., Adams, C., et al. (2016). Association of MTOR Mutations With Developmental Brain Disorders, Including Megalencephaly, Focal Cortical Dysplasia, and Pigmentary Mosaicism. JAMA Neurol 73, 836–845. 10.1001/jamaneurol.2016.0363.

24. Ribierre, T., Deleuze, C., Bacq, A., Baldassari, S., Marsan, E., Chipaux, M., Muraca, G., Roussel, D., Navarro, V., Leguern, E., et al. (2018). Second-hit mosaic mutation in mTORC1 repressor DEPDC5 causes focal cortical dysplasia-associated epilepsy. J. Clin. Invest. 128, 2452–2458. 10.1172/jci99384.

25. Lee, W.S., Stephenson, S.E.M., Howell, K.B., Pope, K., Gillies, G., Wray, A., Maixner, W., Mandelstam, S.A., Berkovic, S.F., Scheffer, I.E., et al. (2019). Second-hit DEPDC5 mutation is limited to dysmorphic neurons in cortical dysplasia type IIA. Annals of clinical and translational neurology 6, 1338–1344. 10.1002/acn3.50815.

26. Yoon, S.J., Elahi, L.S., Pasca, A.M., Marton, R.M., Gordon, A., Revah, O., Miura, Y., Walczak, E.M., Holdgate, G.M., Fan, H.C., et al. (2019). Reliability of human cortical organoid generation. Nat Methods 16, 75–78. 10.1038/s41592-018-0255-0.

27. Uzquiano, A., Kedaigle, A.J., Pigoni, M., Paulsen, B., Adiconis, X., Kim, K., Faits, T., Nagaraja, S., Anton-Bolanos, N., Gerhardinger, C., et al. (2022). Proper acquisition of cell class identity in organoids allows definition of fate specification programs of the human cerebral cortex. Cell 185, 3770–3788 e3727. 10.1016/j.cell.2022.09.010.

28. Trujillo, C.A., Gao, R., Negraes, P.D., Gu, J., Buchanan, J., Preissl, S., Wang, A., Wu, W., Haddad, G.G., Chaim, I.A., et al. (2019). Complex Oscillatory Waves Emerging from Cortical Organoids Model Early Human Brain Network Development. Cell Stem Cell 25, 558–569 e557. 10.1016/j.stem.2019.08.002.

29. Zhang, T., Liu, T., Mora, N., Guegan, J., Bertrand, M., Contreras, X., Hansen, A.H., Streicher, C., Anderle, M., Danda, N., et al. (2021). Generation of excitatory and inhibitory neurons from common progenitors via Notch signaling in the cerebellum. Cell reports 35, 109208. 10.1016/j.celrep.2021.109208.

30. Surmacz, B., Noisa, P., Risner-Janiczek, J.R., Hui, K., Ungless, M., Cui, W., and Li, M. (2012). DLK1 promotes neurogenesis of human and mouse pluripotent stem cell-derived neural progenitors via modulating Notch and BMP signalling. Stem Cell Rev Rep 8, 459–471. 10.1007/s12015-011-9298-7.

31. Bansod, S., Kageyama, R., and Ohtsuka, T. (2017). Hes5 regulates the transition timing of neurogenesis and gliogenesis in mammalian neocortical development. Development 144, 3156–3167. 10.1242/dev.147256.

32. Koopmans, F., van Nierop, P., Andres-Alonso, M., Byrnes, A., Cijsouw, T., Coba, M.P., Cornelisse, L.N., Farrell, R.J., Goldschmidt, H.L., Howrigan, D.P., et al. (2019). SynGO: An Evidence-Based, Expert-Curated Knowledge Base for the Synapse. Neuron 103, 217–234 e214. 10.1016/j.neuron.2019.05.002.

33. Macnee, M., Perez-Palma, E., Lopez-Rivera, J.A., Ivaniuk, A., May, P., Moller, R.S., and Lal, D. (2023). Data-driven historical characterization of epilepsy-associated genes. European journal of paediatric neurology: EJPN: official journal of the European Paediatric Neurology Society 42, 82–87. 10.1016/j.ejpn.2022.12.005.

34. Oliva, M.K., Bourke, J., Kornienko, D., Mattei, C., Mao, M., Kuanyshbek, A., Ovchinnikov, D., Bryson, A., Karle, T.J., Maljevic, S., and Petrou, S. (2024). Standardizing a method for functional assessment of neural networks in brain organoids. Journal of neuroscience methods 409, 110178. 10.1016/j.jneumeth.2024.110178.

35. Blair, J.D., Hockemeyer, D., and Bateup, H.S. (2018). Genetically engineered human cortical spheroid models of tuberous sclerosis. Nature medicine 24, 1568–1578. 10.1038/s41591-018-0139-y.

36. Bacq, A., Roussel, D., Bonduelle, T., Zagaglia, S., Maletic, M., Ribierre, T., Adle-Biassette, H., Marchal, C., Jennesson, M., An, I., et al. (2022). Cardiac Investigations in Sudden Unexpected Death in DEPDC5-Related Epilepsy. Ann Neurol 91, 101–116. 10.1002/ana.26256.

37. Klofas, L.K., Short, B.P., Zhou, C., and Carson, R.P. (2020). Prevention of premature death and seizures in a Depdc5 mouse epilepsy model through inhibition of mTORC1. Hum Mol Genet 29, 1365–1377. 10.1093/hmg/ddaa068.

38. Yuskaitis, C.J., Jones, B.M., Wolfson, R.L., Super, C.E., Dhamne, S.C., Rotenberg, A., Sabatini, D.M., Sahin, M., and Poduri, A. (2018). A mouse model of DEPDC5-related epilepsy: Neuronal loss of Depdc5 causes dysplastic and ectopic neurons, increased mTOR signaling, and seizure susceptibility. Neurobiol Dis 111, 91–101. 10.1016/j.nbd.2017.12.010.

39. Hughes, J., Dawson, R., Tea, M., McAninch, D., Piltz, S., Jackson, D., Stewart, L., Ricos, M.G., Dibbens, L.M., Harvey, N.L., and Thomas, P. (2017). Knockout of the epilepsy gene Depdc5 in mice causes severe embryonic dysmorphology with hyperactivity of mTORC1 signalling. Scientific reports 7, 12618. 10.1038/s41598-017-12574-2.

40. Marsan, E., Ishida, S., Schramm, A., Weckhuysen, S., Muraca, G., Lecas, S., Liang, N., Treins, C., Pende, M., Roussel, D., et al. (2016). Depdc5 knockout rat: A novel model of mTORopathy. Neurobiology of Disease 89, 180–189. 10.1016/j.nbd.2016.02.010.

41. Weckhuysen, S., Marsan, E., Lambrecq, V., Marchal, C., Morin-Brureau, M., An-Gourfinkel, I., Baulac, M., Fohlen, M., Kallay Zetchi, C., Seeck, M., et al. (2016). Involvement of GATOR complex genes in familial focal epilepsies and focal cortical dysplasia. Epilepsia 57, 994–1003. 10.1111/epi.13391.

42. Sloan, S.A., Andersen, J., Pasca, A.M., Birey, F., and Pasca, S.P. (2018). Generation and assembly of human brain region-specific three-dimensional cultures. Nat Protoc 13, 2062–2085. 10.1038/s41596-018-0032-7.

43. Hao, Y., Hao, S., Andersen-Nissen, E., Mauck, W.M., 3rd, Zheng, S., Butler, A., Lee, M.J., Wilk, A.J., Darby, C., Zager, M., et al. (2021). Integrated analysis of multimodal single-cell data. Cell 184, 3573–3587 e3529. 10.1016/j.cell.2021.04.048.

44. McGinnis, C.S., Murrow, L.M., and Gartner, Z.J. (2019). DoubletFinder: Doublet Detection in Single-Cell RNA Sequencing Data Using Artificial Nearest Neighbors. Cell Syst 8, 329–337 e324. 10.1016/j.cels.2019.03.003.

45. Hodge, R.D., Bakken, T.E., Miller, J.A., Smith, K.A., Barkan, E.R., Graybuck, L.T., Close, J.L., Long, B., Johansen, N., Penn, O., et al. (2019). Conserved cell types with divergent features in human versus mouse cortex. Nature 573, 61–68. 10.1038/s41586-019-1506-7.

46. Aran, D., Looney, A.P., Liu, L., Wu, E., Fong, V., Hsu, A., Chak, S., Naikawadi, R.P., Wolters, P.J., Abate, A.R., et al. (2019). Reference-based analysis of lung single-cell sequencing reveals a transitional profibrotic macrophage. Nat Immunol 20, 163–172. 10.1038/s41590-018-0276-y.

47. Bergen, V., Soldatov, R.A., Kharchenko, P.V., and Theis, F.J. (2021). RNA velocity-current challenges and future perspectives. Mol Syst Biol 17, e10282. 10.15252/msb.202110282.

48. Wu, T., Hu, E., Xu, S., Chen, M., Guo, P., Dai, Z., Feng, T., Zhou, L., Tang, W., Zhan, L., et al. (2021). clusterProfiler 4.0: A universal enrichment tool for interpreting omics data. Innovation (Camb) 2, 100141. 10.1016/j.xinn.2021.100141.

